# Information is asymmetry: spatial relations were encoded by asymmetric mnemonic manifolds

**DOI:** 10.1101/2024.03.13.584855

**Authors:** Longsheng Jiang, Yanlin Zhu, Jia Liu

## Abstract

Substantial evidence suggests that working memory (WM) leverages relational representations to provide flexible support for cognitive functions, a capacity likely derived from the dynamic nature of neural codes in WM. However, how these dynamic codes represent and maintain relations remains unclear. Here, we examined the transformation of neural geometries in the dorsal prefrontal cortex of monkeys performing a visuospatial delayed-match/nonmatch task, where the monkeys were instructed to hold the spatial location of a white square in WM to match it with the spatial location of a subsequent square. We found that the sensory manifold during the square’s presence and the mnemonic manifold after the square’s offset both aligned with the stimulus manifold. However, significant differences emerged between the sensory and mnemonic manifolds, exhibiting little correlation in their neural geometries. Further analysis on the dynamic transformation from the sensory to mnemonic manifold revealed a process of expansion followed by flattening: the asymmetric sensory manifold first expanded into a symmetric neural geometry immediately after the square’s onset offset, which then gradually flattened along dimensions different from those initially expanded, culminating in an asymmetric mnemonic manifold. This dynamic process of reconstruction not only remained its faithfulness to the stimulus geometry but also gained the flexibility to meet task demands. In sum, this transformation from asymmetry to symmetry and back to asymmetry in neural geometry precisely illustrates the dynamics of memory reconstruction, shedding lights on the subjective nature of WM that generates both accurate and illusory representation of the world we lived in.

## Introduction

In his dialogue “Theaetetus,” Plato likened memory to a wax tablet, on which our perceptions and thoughts are engraved. This metaphor illustrates how memories are formed and retained, or sometimes forgotten, contingent on the quality of one’s “wax.” This ancient metaphor finds resonance in modern studies of neural geometry embedded in the high-dimensional neural space constructed by the population of neurons. For example, research on the working memory (WM) of monkeys has shown that the neural geometry aligns precisely with the stimulus geometry (Spaak et al., 2017; Xie et al., 2022), reflecting the faithful nature of memory (Murray et al., 2017). However, instances of false memories, where something is remembered differently from its actual occurrence or is recalled as happening when it did not (Loftus, 1993; Loftus & Palmer, 1974), challenge the static nature of neural geometry (Fusi, 2008; Lundqvist et al., 2018; Stokes et al., 2013). The flexibility of memory, while potentially compromising faithfulness, also fosters creativity and problem-solving, supports adaptative functions, and aids in integrating new information with existing memory. An interesting question arises from how the “wax tablet” can simultaneously maintain both accuracy and flexibility in WM.

Several studies have attempted to address this issue (Murray et al., 2017; Cavanagh et al., 2018), proposing that the representations in WM undergo substantial transformations through dynamic coding (Constantinidis et al., 2018; Lundqvist et al., 2018). A prominent phenomenon is that population averages of neuronal activities ramp up over the delay period between stimulus presentation and animal responses, especially in the late delay period (Barak et al., 2010; Rainer & Miller, 2002; Warden & Miller, 2010). Correlated with the up-ramping phenomenon, neural representations reflecting stimulus dissimilarity in a delayed-match/nonmatch task reemerge in the late delay period after being degraded in the early delay period (Barak et al., 2010). The degrading-reemerging of relational representations is also found in the gaze patterns of human participants in a cued-recall WM task (Linde-Domingo & Spitzer, 2023), implying similar changes in the neural representations (Van Ede et al., 2019). Given these results, the late delay period is conceptualized as the information reinstallation phase in WM models (Romo et al., 1999; Stokes, 2015). In this phase, the memorized sensory information is reactivated for supporting later processing. However, how this is realized by neural activities mechanistically is still unclear.

To verify the hypothesis and understand the underlying mechanism, we used a publicly available dataset and analyzed neuronal activities in lateral dorsal prefrontal cortex (PFC) of monkeys, recorded in a visuospatial delayed-match/nonmatch task (Meyer et al., 2011; Qi et al., 2011). This task had two delay periods and only in the first one the monkeys needed to hold the memorandums of the stimuli (see Results). Thus, the first delay period provided an isolated arena for studying relations of spatial stimuli in WM and is the focus of this study. We constructed neural geometries at each moment in the first delay. We found that the resemblance between the neural geometry and stimulus geometry increased monotonically in the late delay period, an indication of reconstruction. The reconstructed neural geometry was different from the neural geometry during stimulus presentation. The reconstruction, on the representation-level, is a process of slightly compressing a symmetric geometry into an asymmetric geometry, while the representation dimensionality remained unreduced. The biased aspect of the asymmetric geometry after compression represents the stimulus relation. The slightly compressed geometry is flexible as it remains high-dimensional, so it retains the possibilities to represent other relations by re-compression, had the task-demand change.

## Results

### Visuospatial delayed-match/nonmatch task and neural geometry analysis

Monkeys were trained to perform a standard match/nonmatch active task (Fig. 1a) (Meyer et al., 2011). In this task, after a 1.0s fixation period, two visual cues of spatial location stimuli were presented serially for 0.5s. Each cue presentation was followed by a 1.5 delay period with no stimuli. The two delay periods served distinct cognitive functions. Demonstrated in previous studies (Meyers et al., 2012) and our decoding analysis (Fig. S1), during the first delay, the monkeys held stimulus information of the first cue in WM, while during the second delay, they made a decision based on the two cues. Then, the monkeys responded through saccade whether the two stimuli were same (matched) or diametrically opposite (non-match). As a control, the monkeys were tested on the passive task, in which the same experimental paradigm was used but the monkey directly received reward without the need of decision-making. In the passive setting, the monkeys were already familiar with the stimuli, though they had not been trained to make any decisions (Meyer et al., 2011).

**Figure 1.**
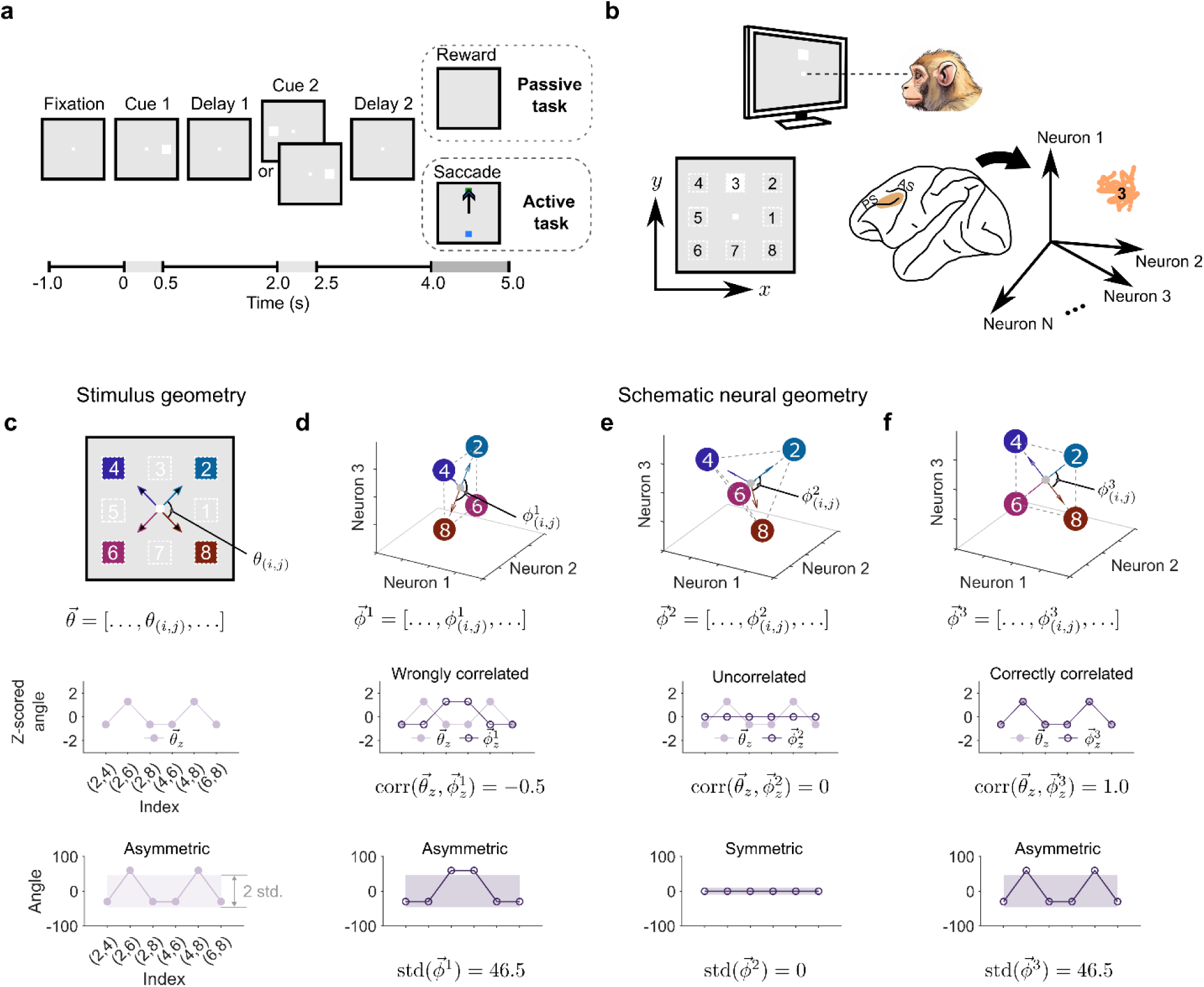
Experiment setup and data analysis methods. **a)** Experiment paradigms of the passive and active tasks. The difference between the tasks was whether the monkeys got reward directly (passive) or needed to make correct match-nonmatch decisions before getting reward (active). **b)** The visual stimuli and the recording site. The monkeys fixated at the center of the 2D screen (top). White squares appeared at one of eight locations (left). Electrophysiological signals of neurons in dorsal lateral prefrontal cortex were recorded and analyzed in a high dimensional neural state-space (right). Neural responses to one stimulus (e.g., stimulus 3) form a point cluster in the space. PS: principal sulcus; AS: arcuate sulcus. **c)** Definition of the angular representational dissimilarity vector (RDV). For illustration, only four of the eight stimuli are depicted in the schematics (top). The z-scored angular RDV shows a signatural pattern (center). Standard deviations of the original angular RDV quantify the degree of geometry asymmetry (bottom). **d-f)** Schematic neural geometries. Definitions of the angular representational dissimilarity vectors of the neural geometries wrongly correlating (d), not correlating (e), and correctly correlating (f) with the stimulus geometry. Shapes of the neural geometries are asymmetric (d), symmetric (e), and asymmetric (f), respectively.

The stimuli were white solid squares appearing on one of nine locations on the monitor screen (Fig. 1b, left). The locations included the fixation point and the surrounding locations evenly distributed on the perimeter of a square. The surrounding eight locations were studied in this work, while the ninth location, the fixation point, was used as the reference. Geometrically, the stimuli located on a 2-dimensional (2D) plane. The spatial relation was established by the ring-shape geometrical layout of the eight possible peripheral locations. Abstractly, the relation could be viewed as a prespecified looped sequence of the stimuli: 1-2-…-8-1. We examined whether and how the relation of the stimuli was represented in WM.

Here, we report the analysis of the physiological recordings of 384 neurons in one monkey (Monkey ELV) participating in both the passive and active tasks. Similar results of two other monkeys, Monkeys ADR and NIN, are in Supplementary. The neurons located in the dorsal lateral prefrontal cortical area, which was over and above the principal sulcus (PS) and adjacent to the arcuate sulcus (AS) (Fig. 1b, right). This region is inherently organized for spatial representations (Meyer et al., 2011). Although most of the neurons were recorded individually, we augmented them together to create a pseudo-population. We analyzed the populational data in a 384D neural state-space (Fig. 1b, right). A neuron’s firing rate at a moment in the task specified the coordinate on an axis. The 384 coordinates determined a neural state which is a point in the neural state-space. The representations of the stimuli in a time interval thus were clusters of the points. We chose cluster centers as the representative of neural representations. These centers together formed a geometrical structure in the neural state-space. We thus could analyze the similarity of the geometrical structure in the neural state-space (neural geometry) and the geometrical layout of the stimuli on the monitor screen (stimulus geometry).

If the neural geometry is similar to the stimulus geometry, the relation among the neural representations matches the relation among the stimuli, and the stimulus relation is therefore represented. Here, we used vectorized representation dissimilarity matrices (representation dissimilarity vectors, RDVs) as the surrogate of a geometry (Fig. 1c-f, top). Specifically, in the stimuli geometry, we drew arrows from the fixation point to the four stimuli. Angle *θ*_(i,j)_ is subtended by the two arrows pointing to stimuli *i* and *j*. The values of the ordered list of *θ*_(i,j)_for any *i* < *j* constructed an angular RDV 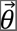 (Fig. 1c). For simplicity, we used four spatial locations and assumed they resided in 3D neural state-spaces (Fig. 1d–f). The angular RDVs 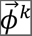 *k* = 1, 2, 3, of the neural geometries were computed with the arrows connecting neural representations of the fixation point and four stimuli. Correlation between the z-scored RDVs 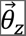 and 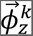 quantifies similarity between the stimulus and neural geometries. The three schematic neural geometries either encode wrong relation (negative correlation, Fig. 1d), no relation (zero correlation, Fig. 1e), or correct relation (positive correlation, Fig. 1f). The correlation acted as the measure of how well the stimulus relation was represented.

Importantly, these distinct correlations further linked to an important characteristic of the neural geometries: symmetry. A geometry is defined as symmetric if it has homogeneous angles and edge lengths. The tetrahedron in Fig. 1e, which is the neural geometry uncorrelated to the stimulus geometry, is an example of symmetric structure, demonstrated by its uniformly distributed angular RDV. In contrast, the 2D stimulus geometry itself is asymmetric, since its angular RDV is not uniform. The degree of asymmetry is described as the extent of deviation of the original RDVs 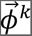 from a uniform distribution, and thus can be quantified by its standard deviation (Fig. 1c, bottom), with larger standard deviations meaning more asymmetric. Accordingly, the correlated neural geometries, whether correctly or wrongly, collapse into the same dimensionality as the stimulus geometry and are asymmetric (Fig. 1d & f). We therefore conjectured that asymmetry of neural geometries might be a necessary condition for representing relations and examined this conjecture with the recorded neural data.

### Simultaneous reemergence of correlation and asymmetry

We first examined how the stimulus relation was maintained in the delay 1 period. We used a sliding window analysis to calculate the correlation between the stimulus and neural geometries over time (−1–2s), including the fixation, cue 1, and delay 1 period (see Methods). The data collected during the active task showed a significant increase in the correlation when the trials entered the cue 1 period, in comparison to the near zero correlation during the fixation period (Fig. 2a). This is expected, as the neural activities were driven by the stimulus presentation, thus termed as the sensory manifold. Interestingly, despite the monkey maintaining the stimulus information, as evident by behaviors in the active task, the correlation quickly fell and reached approximately zero around the cue offset. Then, the correlation ramped up gradually over the mid-late delay 1 period, aligning with the previous findings of the degrading-reemerging phenomenon (Barak et al., 2010). Visualizations of the z-scored angular RDVs in three sampled time windows during the cue 1, early delay 1, and late delay 1 periods confirmed this trend in correlation dynamics (Fig. 2e, right). The neural RDVs in the cue 1 and late delay 1 periods matched the stimulus RDV (Pearson’s *r* > 0.62), but not in the early delay 1 period (Pearson’s *r* = 0.18). This result was further evaluated against baselines.

**Figure 2.**
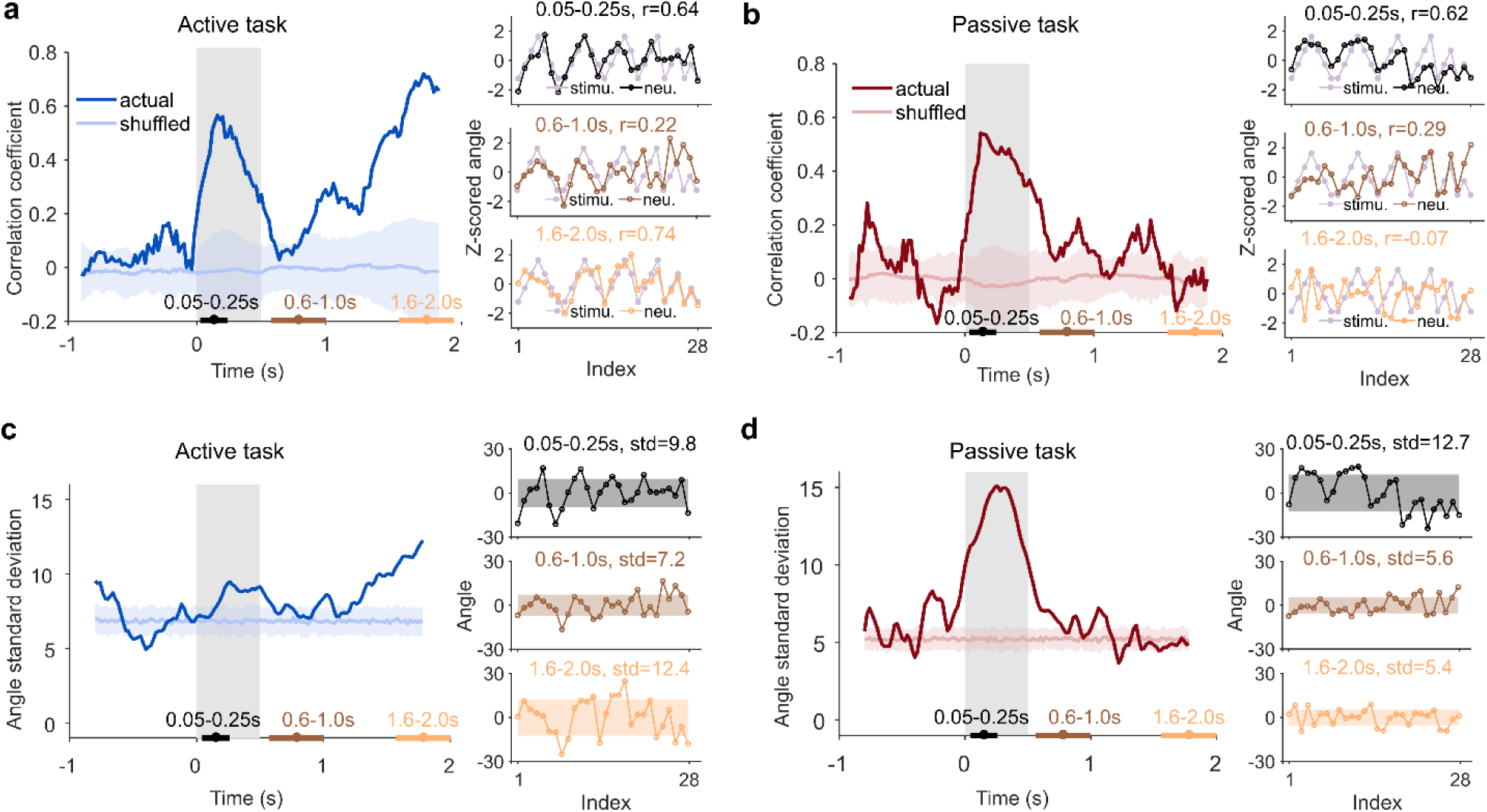
Dynamic correlation and asymmetry of the neural geometries in the active task. **a)** Left, the dynamic correlation between the stimulus and neural geometries in the active task. Randomly shuffling the looped sequence of the stimuli generated the baseline with the means (line) and standard deviations (shadow) as shown. The bottom bars indicate three sampled time windows in the cue 1, early delay 1, and late delay 1 periods, respectively. Right, angular RDVs of the neural and stimulus geometries in the three sampled time windows. r: Pearson’s correlation. **b)** Same as (a) except it is for the passive task. **c)** Left, the dynamic asymmetry of the neural geometries in terms of RDV standard deviation in the active task. The baseline was created by sampling geometries from a space generated by adding Gaussian noise to vertices of the perfectly symmetric geometry. The noise level was adjusted to match the average RDV standard deviation in the fixation period. Right, angular RDVs of the sampled neural geometries and their standard deviations (shadow). std: RDV standard deviation. **d)** Same as (c) except it is for the passive task.

To create a baseline, we randomly shuffled the looped sequence of the stimulus (see Methods) and computed the correlation time courses between the shuffled stimulus geometry and the neural geometries. The averaged correlation of the baseline was approximately zero over the analyzed time interval. The correlations of the active task data in late delay 1 period (1.5–2s) were significantly greater than the baseline (one-tail t-test, averaged *p* < 0.001). A further examination of the correlation between the neural geometry and the permuted stimulus geometries revealed that the correlation monotonically decreased as the dissimilarity between the permuted and actual stimulus geometries increased (Fig. S6). It was the actual stimulus relation that was represented in the delay 1 period of the active task, thus termed as the mnemonic manifold. Similar results were obtained when using a different index of dissimilarity—RDVs of Euclidean distances between neural representations (Fig. S2)—and for the other two monkeys (Fig. S14).

We reasoned that the stimulus relation was represented in the first delay because it was functionally relevant for completing the active task. If it became no longer task-relevant, either in the passive task, where no memorization was required, or in the second delay, where only the match/nonmatch decision was task-relevant, we would expect no relational representation. Indeed, this was true. In the passive task, after the initial surge in the cue 1 due to sensory stimulation, the correlation decreased to and oscillated around zero in the delay 1 period, and became not significantly greater than the baseline in the late delay 1 period (1.5–2s, one-tail t-test, averaged *p* = 0.587, Fig. 2b). This correlation time course was in line with the visible alignment between the RDV of the stimulus geometry and that of the neural geometry sampled in the cue 1 period (Pearson’s *r* = 0.62), and their unalignment in the early delay 1 (Pearson’s *r* = 0.29) and late delay 1 periods (Pearson’s *r* = −0.06, Fig. 2b, right). Similarly, in the delay 2 period of the active task, the correlation oscillated around zero (Fig. S3), showing that the stimulus relation was not represented when stimulus information was task-irrelevant.

Motivated by the conjectured relationship between the stimulus-activity correlation and asymmetry of neural geometries, we next examined the standard deviations of the angular RDVs using a sliding window analysis. Similar as the correlation, in the active task, the RDV standard deviation experienced fall and raise across the cue 1 and delay 1 period (Fig. 1c, left). This trend was confirmed by visualizing the sampled angular RDVs, showing that the RDV standard deviations in the cue 1 (*std* = 9.8) and late delay (*std* = 12.4) were greater than in the early delay (*std* = 7.2) (Fig. 2c, right). On the contrary, the RDV standard deviation in the passive task only experienced a surge in the cue 1 period and then fell back to the same level as in the fixation period (Fig. 2d, left). This trend again was confirmed by visualizing the angular RDVs in the three sampling windows, with RDV standard deviation being large only in the cue 1 period (*std* = 12.7 versus *std* ≤ 5.6) (Fig. 2d, right). To further contrast the temporal changes of the geometry asymmetry, we created a baseline by sampling virtual geometries from a space generated by adding Gaussian noise to a perfectly symmetric geometry. The noise level was adjusted to match the averaged RDV standard deviations in the fixation periods of the active and passive task, respectively. (See Methods). Comparing with the baselines, the actual RDV standard deviations in the late delay 1 period were significantly greater in the active task (1.5–2s, one-tail t-test, averaged *p* < 0.001, Fig. 2c) but not different from the baseline in the passive task (1.5–2s, one-tail t-test, averaged *p* = 0.96, Fig. 2d). The observed reemergence of both the correlation and the asymmetry in the delay 1 period of the active task is in line with our conjecture.

In sum, we verified that the monkeys indeed held the stimulus relation information in the delay 1 period of the active task. However, the relation information was not maintained in a stable code; it underwent a degrading and reemerging process, as if the relational representation was completely reconstructed. Meanwhile, the asymmetry of the neural geometry experienced the same degrading-reemerging trend, suggesting that neural geometries might dynamically represent the stimulus relation through changing their asymmetry. To test whether this is the case, we next explored how the neural geometry transformed during the degrading-reemerging trend.

### Transformation of high-dimensional neural geometry supports relation reconstruction

Presumably, WM maintains a piece of memory by caching an exact copy of its sensed information. However, the degrading-reemerging trend in the stimulus-activity correlation disagrees with this intuition and suggests a more complex reconstructive mechanism instead. If the stimulus relation was completely reconstructed, its reconstructed representation in WM would likely be distinct to the relational representation driven by sensing. To see whether this was true, we applied PCA to the neural representations sampled in the cue 1, early delay 1, and late delay 1 periods. We found in the cue 1 and late delay 1 periods, the projections of the neural representations to the first two principal components (PCs) both had a looped sequence structure (Fig. 3a, left & right). The structures were consistent to the stimulus geometry. They were in contrast to the neural geometries sampled in the early delay 1 period (Fig. 3a, middle) or across the whole delay 1 period in the passive task (Fig. S4 middle & right), where the projections appeared rather random. Importantly, the looped structures in the cue 1 and late delay 1 periods, respectively, were different in their exact shapes: while the cue 1 structure elongated more along the line connecting representations of stimulus 1 and 6, the late delay 1 structure elongated less along this line. This observation supports the reconstructive view of WM.

**Figure 3:**
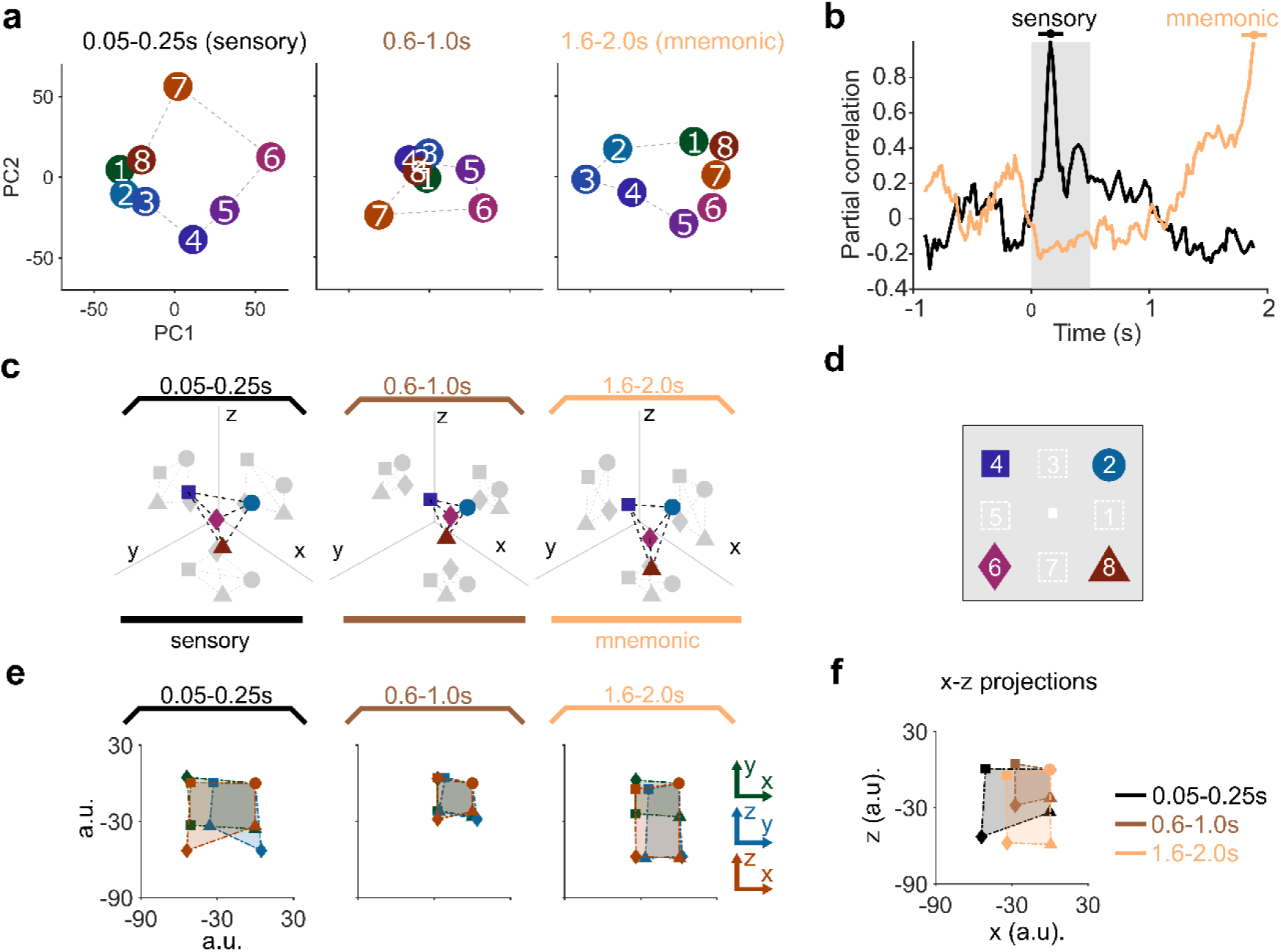
Visualization of the reconstructive neural geometries. **a)** Projections to the first two PCs of the neural geometries of the active task data in the three sampling time windows. **b)** The cross-temporal partial correlation with respect to the neural geometries in the sensory and mnemonic windows, respectively, while controlling for the effect of the stimulus geometry in the active task. **c)** Visualization in 3D spaces of the neural geometries formed by representations of the stimuli in (d) and in the three sampled time windows in the cue 1, early delay 1, and late delay 1 periods, respectively. Marker shapes refer to stimulus identities. **d)** The stimulus geometry with four selected spatial stimuli. **e)** The 2D projections of the neural geometries in (c), grouped according to the sampling time windows. **f)** The *x*-*z* projections in the three sampling time windows.

To quantify the difference between the sensed and reconstructed relational representations, we used partial correlations between the neural geometries while controlling the effect of the stimulus geometry. Specifically, we denoted the neural geometry sampled in the cue 1 period as the sensory manifold and that in the late delay 1 period the mnemonic manifold. We computed cross-temporal partial correlation of the neural geometries across time with reference to the sensory or mnemonic manifold, respectively (Fig. 3b). As the neural geometry moved away from the sensory manifold, its partial correlation to the sensory manifold decreased from 1 and approached −0.2 when it arrived at the mnemonic manifold. Reversed trend was observed in its partial correlation to the mnemonic manifold. The sensory and mnemonic geometries were indeed different; the mnemonic manifold was actively reconstructed. Despite their difference, both the sensory and mnemonic geometries could prioritize the relational representation to the first 2 PCs. These results suggest that the purpose of reconstruction was not to create an exact copy of what was sensed but to only prioritize the relation, which required much less cognitive effort. We were interested in how the prioritization was enabled by the transformation of the neural geometry, and how this transformation led to the change in its asymmetry.

For gaining insights, we started by visualizing the neural geometries directly in a 3D space. To do so, we focusing on four stimuli’s representations in the active task, stimulus 2, 4, 6, and 8 (Fig. 3c), before extending analysis to the full set of stimuli. These stimulus representations created a neural geometry similar to the schematic neural geometries above (Fig. 1d–f). The axes of the 3D space were obtained with linear support vector machine (SVM). Specifically, we classified representations of the stimuli {2, 8} versus {4, 6}, {2, 6} versus {4, 8}, and {2, 4} versus {6, 8} with linear SVM. The optimal separation vector was the tentative *x*-, *y*-and *z*-axis, respectively. The three tentative axes were orthogonalized by the Gram-Schmits method (Strang, 2022), creating a 3D Euclidean space. Neural representations were then projected to this 3D space.

We first visualized the sensory manifold in the cue 1 period (0.05–0.25s) where the stimulus relation information was encoded (Fig. 3c, left). The stimulus relation was not represented as we speculated in the schematic neural geometry above (Fig. 1f), where the neural geometry collapsed into the same dimensionality as the stimulus geometry. Instead, the sensory manifold took the highest dimensionality (3D) afforded by four representations. Crucially, its shape was asymmetric, consistent with the asymmetry measure in Fig. 2c. More specifically, its projection to the *x*-*z* plane was greater than that to the *x*-*y* and *y*-*z* plane (Fig. 3e, left, average ranked area ratio = 1.27, see Methods). The 2D *x*-*z* projection had the same looped sequence as the stimulus geometry (Fig. 3d). The larger area encompassed by the *x*-*z* projection made this particular looped sequence stand out among other possible looped sequences, thus prioritizing the information of the stimulus relation in the high-dimensional sensory manifold.

We next visualized the neural geometry in the early delay 1 period (0.6–1.0s) where the correlation between the stimulus and neural geometries was low. We found the neural geometry was nearly symmetric (Fig. 3d, middle) for its three 2D projections were almost identical (Fig. 3e, middle, average ranked area ratio = 1.05). This is consistent with the symmetric schematic neural geometry that is of zero correlation to the stimulus geometry (Fig. 1e). As a result, no relational information was prioritized in this neural geometry and no relation was represented. On the other hand, we visualized the neural geometries sampled in the delay 1 period in the passive task where it was known that relational information was not memorized. The neural geometries in this period were also symmetric (Fig. S8). Therefore, the stimulus relation was not represented if and only if the neural geometry were symmetric. The symmetric neural geometry provided a neutral reference against which the sensory and mnemonic geometries could be examined.

Similar to the sensory manifold, the mnemonic manifold in the late delay 1 period (1.6–2.0s) was high-dimensional and had an asymmetric shape (Fig. 3c, right). Again, the *x*-*z* projection was the largest among all 2D projections (Fig. 3e, right, average ranked area ratio = 1.69) [stats], showing that the stimulus relation was prioritized. Nevertheless, the enlarged *x*-*z* projection of the mnemonic manifold was different than that of the sensory manifold (Fig. 3f), consistent with the partial correlation analysis in Fig. 3b, visually justifying that the mnemonic manifold was indeed reconstructed anew. Taken together, the neural geometries sampled in the three discrete time windows hint a representation-level mechanism for representing relations in WM: During the delay 1 period, symmetric high-dimensional neural geometries actively deform to asymmetric ones for prioritizing the stimulus relation. We next investigated this idea with the neural geometries formed by all the eight representations in the active task.

### From symmetry to asymmetry: targeted flattening of neural geometries

To investigate the process from the symmetrical neutral reference to the asymmetric mnemonic manifold, we analyzed the neural data across the mid-late delay 1 period (1-2s). In this interval the stimulus-neural geometry correlation increased monotonically. By definition, when applying PCA to symmetrical neural geometries, the explained variances are uniformly distributed. As the degree of asymmetry increases, the uniformity decreases. We therefore applied PCA to the neural data to evaluate asymmetry in a series of 0.2s-wide time windows for both the passive and active tasks (Fig. 4a). Of the passive task data, the explained variances were initially close to uniform distribution, reflecting a symmetrical neural geometry. This is consistent with the above observation that symmetrical neural geometries did not represent relations. The explained variances remained uniform and constant over time, as we would expect for the passive task data. The explained variances of the active task data initially were also close to uniform distribution. However, as time proceeded, the explained variance distribution experienced a monotonic decrease in uniformity. Specifically, the explained variances of PC 1 and PC 2, especially of PC 1, continuously increased, while that of other the PCs remained approximately constant. It implied that of the active task data the breakdown of symmetry in the neural geometries was gradual and continuous. In sum, the transformation from symmetry to asymmetry was a monotonic process.

**Figure 4:**
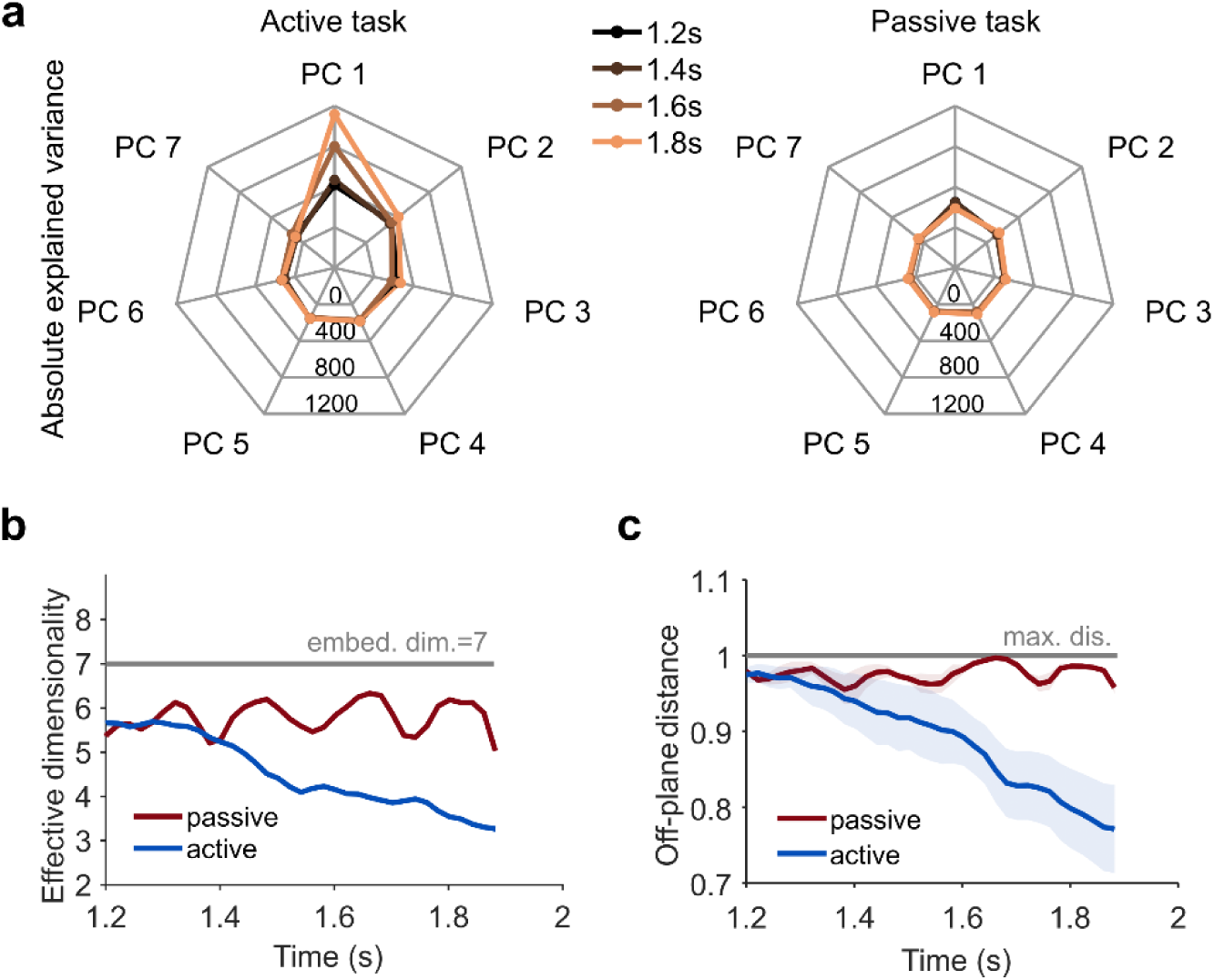
Transformation from a symmetric neural geometry to an asymmetric neural geometry. **a)** The absolute explained variances of the PCs over the mid-late delay 1 period in the active (left) and passive (right) tasks. **b)** The effective dimensionality of the neural geometry over the mid-late delay 1 period in the active and passive tasks. Gray line: the embedding dimensionality. **c)** Similar to (b) but showing the averaged off-plane distance. Shading areas: the standard deviation. Gray line: the upper bound of the off-plane distance.

Given dimensionality of neural geometries is important in measuring captured task information (Chung & Abbott, 2021), we examined the effect of the symmetry-to-asymmetry process on dimensionality. We first analyzed embedded dimensionality which is the minimal number of Euclidean dimensions required for containing neural geometries and is relevant to how information is processed (Jazayeri & Ostojic, 2021). Embedded dimensionality can be estimated from the number of linearly separable binary classifications (Rigotti et al., 2013). In this case, the embedded dimensionality was constantly 7D across time (Fig. 4b, gray line), in both the active and passive tasks demonstrated by the actual number of separable classifications of the stimulus representations equaling the theoretical maximum afforded by the eight representations (2^8^ = 256). It is consistent with the above visual observations (Fig. 3c) and the previous studies finding that neural populational coding is high-dimensional (Stringer et al., 2019).

However, the constant embedded dimensionality failed in reflecting the changes afforded by the symmetry-to-asymmetry process. We therefore used a more nuanced index of dimensionality, effective dimensionality, which is a continuous quantity measuring the equivalent number of orthogonal dimensions, each with equal variance (Del Giudice, 2021). We calculated the effective dimensionality for the passive and active task data (See Methods) (Fig. 4b). In the passive task, the effective dimensionality remained relatively constant, despite local fluctuation. Meanwhile, in the active task, the effective dimensionality decreased monotonically from near 7D to approximately 3D, showing clear evidence of dimensionality compression. Since effective dimensionality relates to information entropy (see Methods)—high effective dimensionality means high entropy (Del Giudice, 2021) (Fig. S9)—we would interpret the this dimensionality compression as reducing the uncertainty of the represented information. It implies that the reconstructed the relational representation in this process gradually became more certain. Similar results regarding effective dimensionality were observed in the other two monkeys’ data (Fig. S14).

To further comprehend the process of dimensionality compression in reconstructing the relation of the spatial stimuli in this work, we developed a geometric index to specifically measure how well a neural geometry morphed toward the stimulus geometry. Decreased effective dimensionality is associated with flattened data geometries: explained variances along some dimensions are smaller than along the others (Del Giudice, 2021). We therefore hypothesized that along the dimension compression process, the neural geometry was flattened to approach the stimulus geometry. A way to measure flatness was to evaluate how well in the neural geometry the normalized diametric vectors, defined according to the stimulus geometry, were coplanar (for details, see Method). For instance, in extreme, the neural geometry had been flattened to the 2D stimulus geometry. In this ideal case, all diametric vectors had been completely coplanar. To measure coplanar-ness, we randomly chose two diametric vectors as the basis to span a plane and computed residuals between the other diametric vectors and their projections on the plane. The length of the averaged residual was called off-plane distance. Only when a neural geometry approaches the stimulus geometry in Fig. 1c, does the off-plane distance descend. We analyzed the off-plane distance over the mid-late delay. The averaged off-plane distance of the active task data monotonically decreased from nearly the maximal off-plane distance value to a moderate value, reflecting the transformation to a high-dimensional but flattened geometry (Fig. 4c). In contrast, the averaged off-plane distance of the passive task data remained close to the maximal value. Similar results for other monkeys are in Fig. S14. Taken together, the relational representation was indeed reconstructed through a dimensionality compression process, which flattened the symmetric neural geometry towards the stimulus geometry.

## Discussion

Recent studies reveal that working memory (WM) is much more complex and dynamic than previously thought (Buschman & Miller, 2023; Stokes et al., 2013) and modelled (Baddeley, 2010; D’Esposito & Postle, 2015). What neural substrates encode memorized content, and how to reconcile dynamic coding and stable memoranda have been a heavily debated topic (Cavanagh et al., 2018; Constantinidis et al., 2018; Lundqvist et al., 2018; Miller et al., 2018; Mongillo et al., 2008; Murray et al., 2017). Here we investigated how spatial relations were represented and memorized in dynamic WM. We followed the neural population doctrine (Ebitz & Hayden, 2021; Libedinsky, 2023; Saxena & Cunningham, 2019) and analyzed a pseudo-population data of neurons recorded in a visuospatial delayed-match/nonmatch task (Meyer et al., 2011; Qi et al., 2011), aiming to provide a mechanism for relational representations in WM. We found that during the sensory phase and memory phase the neural geometries morphed to geometries highly correlated with the stimulus geometry. The active deformation of neural geometries is compatible with dynamic coding in WM. Crucially, in the late-delay 1 period, where task-relevant information needed to be recalled, we found the deformation realized progressive reconstruction of the stimulus relation, serving the purpose of information reinstallation (Stokes, 2015). Further, we revealed the relation reconstruction arose from a process transforming symmetric geometries to asymmetric geometries. This transformation monotonically compressed the effective dimensionality, leading the aspect reflecting the stimulus geometry being dominant. Taken together, these results show that relation information is represented in neural geometries, and maintenance of the information in WM is a dynamic reconstructive process, taking place by transforming to asymmetric neural geometries. This mechanism of WM has the potential to be extended to other more semantic relations, and it provides a new perspective helping settle the ongoing stable versus dynamic coding debate.

Consistent with previous works (Barak et al., 2010; Bouchacourt & Buschman, 2019; Murray et al., 2017), the sensory and mnemonic geometries were less correlated, indicating that the mnemonic manifold was created anew. It is peculiar that from the sensory manifold to the mnemonic manifold, there was an intermediate phase, in the early delay, where the stimulus relation degraded. The stimulus relation was then reconstructed during the mid-late delay 1 period. This degrading-reemerging trend was evident in previous works for the correlation between neural population coding of neighboring vibration frequencies (Barak et al., 2010); for the accuracy of stimulus decoding via dynamic coding subspaces (Murray et al., 2017); and for the correlation between the gaze pattern and the stimulus geometry (Linde-Domingo & Spitzer, 2023). This reconstructive view of WM is in line with the studies investigating response biases in visual WM (Poirier et al., 2017; Scotti et al., 2021) and coincides with the notion of reconstructive memory in general (Roediger & DeSoto, 2015; Xue, 2022). During reconstruction, sensed information integrates with internal knowledge, rendering subjective, rather than objective, representations, and occasionally leads to false memory (Mendez & Fras, 2011). Dynamic codes in WM therefore serve the important purpose of reconstruction.

In an effort to reconcile dynamic coding and stable memoranda in WM, several works proposed a theory claiming that there exist stable subspaces in dynamic codes (Murray et al., 2017; Sapountzis et al., 2022; Spaak et al., 2017). In these stable subspaces, stimulus relations are always present, against the reconstructive process found in this work. This discrepancy can be explained by different task demands required in these and in our works (Cueva et al., 2020). Different tasks used in studies require either prospective or retrospective codes in the delay period (Purg et al., 2022). Tasks in which stable subspaces were detected have straightforward stimulus-response mapping, for example, the oculomotor delayed response task (Murray et al., 2017), memory-guided saccade (Spaak et al., 2017), and the spatial attentional cue task (Sapountzis et al., 2022). Therefore, the subjects used prospective coding (Rainer et al., 1999): they immediately plan motor responses along sensory encoding and maintain the stable response information over the delay (Boettcher et al., 2021). Meanwhile, tasks in which reconstructive process were detected have no clear stimulus-response mapping, for example, the vibrotactile delayed discrimination task (Murray et al., 2017), the duration discrimination task (Genovesio et al., 2009), and the color attentional cue task (Sapountzis et al., 2022). These tasks are more complex than, for instance, the memory-guided saccade task (Spaak et al., 2017), and reactivating sensory information before responding becomes necessary (Purg et al., 2022). The reason why reconstruction is necessary lies in the characteristics of each task.

The visuospatial delayed-match/nonmatch task used in this work had two task-relevant cues presented in sequence before decision-making. Correct decisions relied on non-interfering neural representations of the memorized cue and of the sensed cue at the cue 2 period, respectively (Fig. 1a). The neural space likely used orthogonal subspaces to respectively store sensory and memory information (Gurnani & Cayco Gajic, 2023). Thus, the same sensory subspace was used for representing both cue 1 and cue 2. This is true since a linear decoder, trained on data in the cue 1 period, generalized well to the data in the cue 2 period, but poorly in the delay 1 periods (Fig. S14). To avoid interference, after representing cue 1, the sensory subspace needed to be emptied, before it was reused for representing cue 2. The cue 1 representation transported to the orthogonal memory subspace through, for instance, rotational dynamics (Libby & Buschman, 2021), showing higher degree of dynamicity around the cue offset (Sapountzis et al., 2022). This complex transient relocation made the represented relational information temporarily lost. Taken together, the reconstructive process identified in this work calls into question the stable subspace theory as the universal solution to reconciling the dynamic and stable coding of WM.

Through the lens of neural geometries, the relation reconstruction was realized by transforming a symmetric geometry to an asymmetric geometry. At first glance, it may seem counterintuitive that the brain forms symmetric neural geometries when it is in idle, for instance, in the delay 1 period in the passive task, contrasting our intuition that since Plato symmetry has been linked to beauty and is effortful to get in our 3D world (Saller, 2017). However, due to mathematical properties, in high-dimensional spaces, such as the neural space formed by hundreds of neurons, symmetrical geometries are the easiest to obtain, for instance, through random embedding (Gorban et al., 2020). The focus then should be on how asymmetric geometries realizing cognitive functions (Bernardi et al., 2020; Wójcik et al., 2023), and in this work, encoding relational information in the mnemonic manifold.

Our results show the asymmetric mnemonic manifold was transformed from a symmetric geometry. This transformation could be quantified by the descending effective dimensionality. The net effect was compressing the relative variances of the task-irrelevant axes while enlarging that of the task-relevant axes, making the neural geometry flat. Along the enlarged axes, the neural geometry matched the stimulus geometry, thus representing the stimulus relation. Then, the downstream regions might use Hebbian-type learning to implement PCA algorithms (Oja, 1982, 1989) for reading out the relational information. This read-out strategy is more flexible than the read-out assumption of a static classifier (Parthasarathy et al., 2019; Spaak et al., 2017), as it does not necessitate stable subspaces in the neural activities. Further, the PCA read-out strategy does not require the neural geometries to be compressed completely to the 2D stimulus geometry. As long as the variance asymmetry becomes enough for the stimulus relation to be picked up by PCA, the relation information can be successfully read-out without further compression. Our results are in support of this conjecture, the dimensionality of the neural geometries at any moment remained high-dimensional in terms of embedding dimensionality. The high dimensionality, furthermore, offers additional flexibility to the neural representations. Without hard dimension reduction, no feature information was completely lost (Flesch et al., 2022; Grand et al., 2022; Ma et al., 2023). The retained feature information could readily serve different demands, had the task changed. Taken together, the symmetry-to-asymmetry process suggests a mechanism for realizing cognitive flexibility in dynamic WM (Stokes et al., 2013). That is, flattening a high-dimensional neural geometry that is rich in information (Flesch et al., 2022; Sreenivasan et al., 2014) to biasing task-relevant information only, and implementing PCA algorithms in downstream for read-out. Once the current task-demand is lifted, the neural geometry recover to symmetry, ready for new tasks. This mechanism has the potential to be extended beyond spatial relations to semantic relations.

In sum, this work proposed a neural geometry-based mechanism to describe the dynamic process of relation reconstruction in the lateral-dorsal PFC area. The reconstruction was achieved through flattening symmetric neural geometries to asymmetric ones. Asymmetric neural geometries have been found in other brain areas and serving other cognitive functions (Bernardi et al., 2020). Whether the proposed mechanism is universal across brain areas deserves independent future studies. Nevertheless, the mechanism is only on representation-level; its implementational details in terms of neural circuits require further in-depth investigation (Mongillo et al., 2008; Stokes, 2015; Szatmáry & Izhikevich, 2010). Still, even the representation-level mechanism provides us a new venue to explore the causes of relational representation and more cognitive functions across the brain.

## Methods

### Experimental design and neural dataset

This study reanalyzed neurophysiological data from previous investigations involving rhesus monkeys (Macaca mulatta) performing a visuospatial delayed-match/nonmatch task (Constantinidis et al., 2016; Meyer et al., 2011; Qi et al., 2011; Tang et al., 2022). Monkeys sat in a primate chair, fixating on a 0.2°white square while 2°wide white squares appeared randomly in a 3 ×3 grid. The experiment had passive viewing and active decision-making phases, separated by months of training. Liquid rewards were given for maintaining fixation (passive phase) or correctly executing saccades to green or blue targets (active phase). Error trials were limited, with analysis focusing on correct responses (8% error rate).

Recording sites were in the dorsolateral prefrontal cortex, covering the principal sulcus and extending posterior to the arcuate sulcus, including area 46 and parts of areas 8a. The dataset comprised 384 neurons from monkey ELV during both tasks. The raw neuronal data were converted to firing rates for analysis (Constantinidis et al., 2016; Meyer et al., 2011; Qi et al., 2011; Tang et al., 2022). The raw data for each neuron consisted of spike trains, representing a time series detailing neural firing peaks within a trial.

To convert a spike train into a peristimulus time histogram (PSTH), a Gaussian kernel with a standard deviation of 0.01 was applied, sliding across the spike train at a sampling frequency of 500Hz.

For each stimulus and neuron, the PSTHs of all trials, encompassing both matched and nonmatched cues, were averaged. This trial-averaged PSTH served as the neuron’s firing rate in response to the stimulus. More detailed description, see Supplementary Methods M.1.

### Stimulus-neural geometry correlation

Our analysis focused on populational data within a 384-dimensional neural state-space. Each neuron’s firing rate at a given moment defined a point in this state-space. Within a brief time window, a stimulus activated a cluster of points, with the cluster center serving as the neural representation of the stimulus. The neural representations of various stimuli collectively constituted a geometrical structure termed as a neural geometry. Characteristics of this neural geometry were summarized into representational dissimilarity vectors (RDVs). We predominantly utilized angular RDVs. An angular RDV comprised the angles between any two of the ordered arrows emitting from stimulus 9 (or its neural representation) to one in stimuli 1 to 8 (or its neural representation) (Fig. 1c–f). Accordingly, each angular RDV had 28 angular values.

To quantify dynamic similarity between the stimulus geometry and neural geometries, we employed the correlation between RDVs in a sliding time window (width: 0.2s; time step: 0.02s). To enhance robustness, we divided the window into 4 sub-windows, generating 28-component RDVs for neural geometries in each sub-window. Z-scoring eliminated misleading correlations caused by magnitude differences across sub-windows. The z-scored RDVs were concatenated into a 112-component pattern. Simultaneously, the 28-component RDV of the stimulus geometry was repeated 4 times, z-scored, and concatenated into a 112-component pattern. Pearson’s correlation coefficient was then computed between these patterns, providing stability and reliable reflection of actual geometry similarity.

To assess the similarity between sensory and mnemonic geometries, we employed partial correlation to mitigate the impact of the stimulus geometry. The sensory geometry was defined as the neural geometry formed by neural representations within the 0.05–0.25s window, while the mnemonic geometry encompassed representations within the 1.6–2.0s window. Using the sensory geometry as the reference, we compared angular RDVs of neural geometries across a trial against that of the reference. Let the Pearson’s correlation between the neural geometry (N) in a time window and the reference geometry (R) be *r_NR_*; between the neural geometry and the stimulus geometry (S) *r_NS_*; between the reference geometry and the stimulus geometry *r_RS_*. The partial correlation between the neural geometry and the reference geometry while controlling the effect of the stimulus geometry is

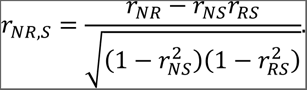

The dynamic partial correlation was computed used a sliding window method whose width was 0.2s and time step was 0.02s. Later, we used the mnemonic geometry as the reference and computed partial correlation in the same way.

### Measuring Neural Geometry Asymmetry

To quantify neural geometry asymmetry, we utilized the standard deviation of angular RDVs, where a larger standard deviation indicated higher asymmetry. We calculated the 28-component RDV for a neural geometry formed by neural representations within a 0.4s-wide time window. Dynamic asymmetry was computed by sliding this time window with a 0.02s time step. For the baseline, we generated a perfectly symmetric geometry by initializing a 384×8 zero matrix and assigning an 8×8 identity matrix to the first 8 rows, representing vertices in the 384-dimensional space. Gaussian noise was added to each dimension of the vertex locations, with noise levels adjusted to match the averaged asymmetry measure during fixation periods. The Gaussian noise was *N*(0, 0.118) for the active task and *N*(0, 0.088) for the passive task. To ensure arbitrariness, the symmetric geometry was randomly rotated in the 384-dimensional space. We sampled 100 geometries, using their angular RDVs to compute the baseline asymmetry.

### Visualization of neural geometries in 3D space and area ratio of 2D projections

Without losing generality, we visualized neural geometries formed by 4 neural representations corresponding to stimuli 2, 4, 6, and 8 in a 3D space defined by the coordinate axes specified using support vector machines (SVM). For the four different stimuli, there were three equal dichotomous classifications that were independent of each other: {2, 4} versus {6, 8}, {2, 6} versus {4, 8}, and {2, 8} versus {4, 6}. We applied linear SVM to each of the classifications and the resulting normalized optimal separation vectors, 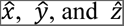, were approximately orthogonal. We then forced the axes to be mutually orthogonal by applying QR decomposition (see more details in Supplementary Methods M.7). After that, the 3D space formed by three mutually orthogonal axes was used to visualize the stimulus representations.

Once the neural geometries formed by 4 neural representations were visualized in a 3D space, we immediately got their 2D projections on the *x*-*y*, *y*-*z*, and *x*-*z* planes. The looped sequence in each 2D projection created a polygon. We used the shoelace formula to compute the area of the polygons: *S_xy_*, *S_yz_*, and *S_xz_*. For instance,

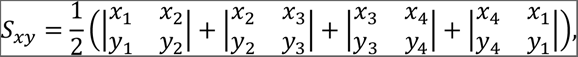

where (*x_i_*, *y_i_*) is the *i*th point of the *x*-*y* projection. We ranked the areas [*S_xy_*, *S_yz_*, *S_xz_*] in a descending order as [*S_hi_, S_mid_, S_lo_*]. The ranked area ratios were

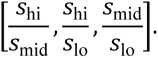

The average ranked area ratio took the mean of the ranked area ratios.

### Effective dimensionality and information entropy

Effective dimensionality (ED) is closely related to the explained variances computed by PCA. For the 8 neural representations in a time window, the ranked explained variances, in descending order, were [λ_1_, λ_2_, …, λ_7_]. The effective dimensionality of the neural representations was defined as

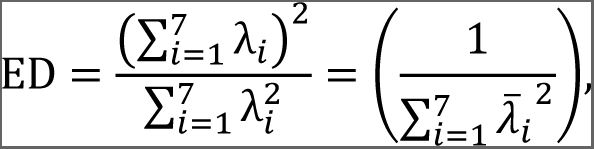

where 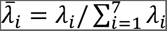 was the normalized explained variance. ED has a connection to Shannon’s information entropy, providing it an information-theoretic interpretation. For the 8 neural representations in our study, the embedded dimensionality was 7, giving rise to 7 feature axes *F_i_*, 1 ≤ *i* ≤ 7. The normalized explained variances along feature axes *F_i_* was 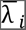, which have the following properties: (1) 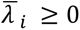 for 1 ≤ *i* ≤ 7; (2) the normalized explained variance of a *n*-D subspace is 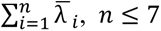; and (3) 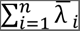= 1. Therefore, the normalize explained variances fulfill the three axioms of probability, thus can be interpreted as the probability of representing feature *F_i_* by the neural geometry: 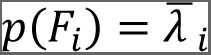. We then computed Shannon’s entropy

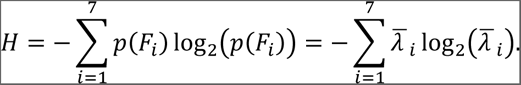

For a complete interpretation of the ED and information entropy, refer to Supplementary Methods M.10.

### Off-plane distance

We introduced the off-plane distance as a measure to assess the extent of flattening in a neural geometry toward a structure resembling the stimulus geometry. We randomly selected two diametric vectors, identified the plane spanned by them, and computed the distance residual between each remaining diametric vector and its projection onto the identified plane. This distance residual, termed the off-plane distance, served as a metric for the flatness of the neural geometry, with a higher averaged off-plane distance indicating a less flat neural geometry. For the reason of this definition, refer to Supplementary Methods M.11.

More methods that were used are reported in detail in Supplementary Methods.

## Notes

### Competing Interest Statement

The authors have declared no competing interest.

