## Supplementary Materials for "Information is asymmetry: spatial relations were encoded by asymmetric mnemonic manifolds"

### Supplementary Methods

In this Supplementary Methods, we present detailed versions of the methods in the main text.

#### M.1. Experiment setup and Neural dataset

We reanalyzed neurophysiological data from previous studies (Constantinidis et al., 2016; Meyer et al., 2011; Qi et al., 2011; Tang et al., 2022), investigating rhesus monkeys (Macaca mulatta) in executing a visuospatial delayed-match/nonmatch task. The detailed description of the task is in (Meyer et al., 2007). Briefly, monkeys sat in a primate chair while a monitor was placed about 60 cm away. The monkeys were required to fixate at a 0.2° wide white square at the monitor center throughout the trial. The spatial stimuli were 2° wide white squares which appeared in one of 9 possible locations in a 3 × 3 grid with 10° spacing between the stimuli. The stimuli appeared randomly across trials. The monkeys were familiarized with all the stimuli in training before the recorded experiment sessions.

The experiment had two phases, the passive viewing (passive task) and active decision-making phases (active task), separated by the several month-long training (Qi et al., 2011; Tang et al., 2022). In the passive phase, the monkeys received liquid reward after successfully maintaining fixation till the end of the second delay. In the active phase, after the second delay period, the monkeys needed to make a saccade to a green target if the two presented stimuli were at the same location; a blue target if they were at diametrically opposite locations. The location of targets varied randomly across trials but were orthogonal to the stimuli. The random location of the green and blue targets prevented the monkeys planning movement direction beforehand (Meyers et al., 2012). Only when the saccade responses were correct could the monkeys receive liquid reward. Since the trials in which monkeys made errors were limited (overall error rate was 8%), only the correct trials were analyzed (Kobak et al., 2016; Qi et al., 2011).

Neurophysiological data from three monkeys, monkey ELV, monkey ADR, and monkey NIN, were used in this study. The data from monkey ELV and monkey ADR were published together with (Kobak et al., 2016). The data from monkey NIN were published together with (Tang et al., 2022). For detailed surgery and neurophysiology information, including anatomical localizations, microelectrode specifications, surgical operations, data acquisition, sorting algorithms, please refer to these studies (Meyer et al., 2011; Qi et al., 2011; Tang et al., 2022). Briefly, the recording sites were in the dorsolateral PFC, which was an area covering the two banks of the principal sulcus (≤2 mm from the center of the principal sulcus) and extending posterior to the arcuate sulcus, incorporating the posterior aspect of area 46 and parts of areas 8a (Meyer et al., 2011; Meyers et al., 2012; Tang et al., 2022). In total, 384 neurons of monkey ELV were recorded in the dataset (Constantinidis et al., 2016) for both the passive and active tasks (Fig. M1a). The number of recorded neurons in Monkey ADR was 137 in the active task, and 65 in the passive task. These neurons were used without further selection. For monkey NIN, since its neurons were recorded in tasks using varying delay lengths (0.35–1.5s), we selected neurons having delay lengths greater than 1s and truncated the delay periods to 1s (Tang et al., 2022). The number of selected neurons of monkey NIN was 141 and 112 in the active and passive tasks, respectively. The raw data of the neurons were then converted to firing rate.

The raw data of each neuron were spike trains, consisting of a time series detailing neural firing peaks in a trial. To transform a spike train in a trial to peristimulus time histogram (PSTH), a Gaussian kernel with a standard deviation of 0.01 was applied, sliding across the spike train. The sampling frequency of applying the Gaussian kernel was set as 500Hz. For each stimulus and each neuron, the PSTH of all trials were averaged, including trials in which the two cues were matched or nonmatched. The trial-average of PSTH served as the firing rate of the neuron in response to the stimulus. The averaged firing rate of all neurons showed clear responses to the cue onsets (Fig. M1b–d), with the averaged firing rate in the passive task lower than that in the active task.

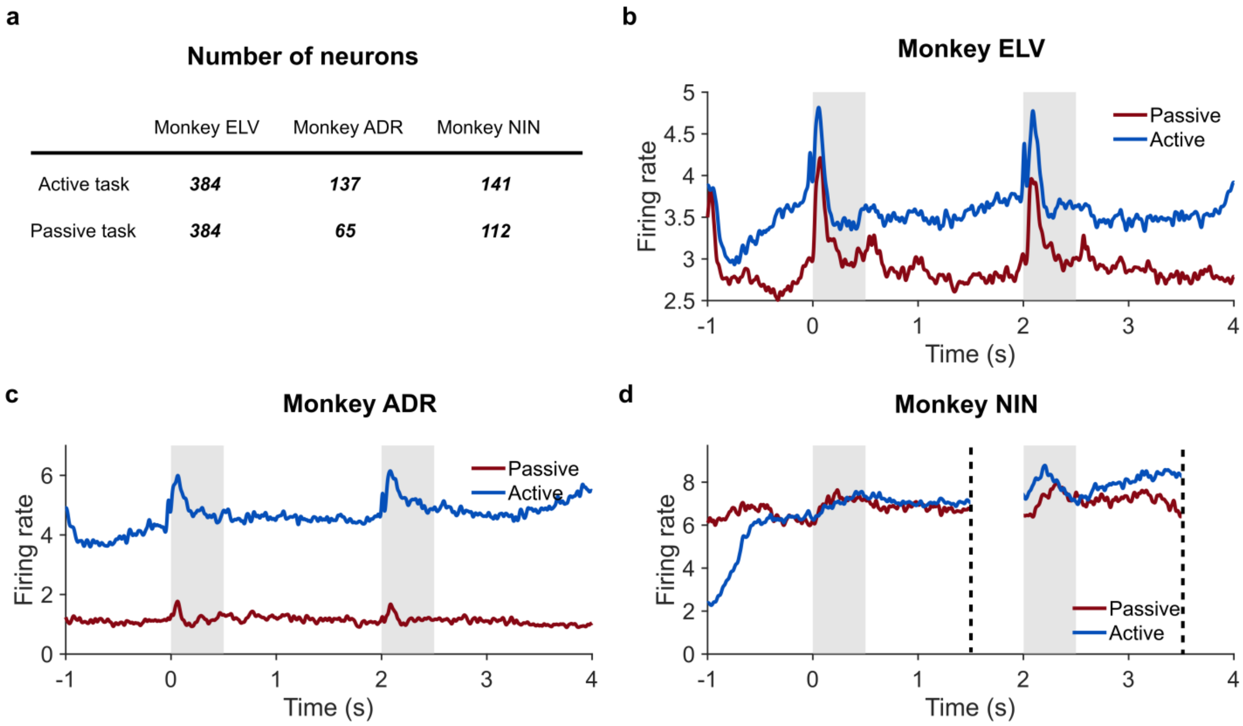


**Figure M1.** Number of the recorded neurons and their average peri-stimulus time histograms. **a)** The numbers of the recorded neurons in the dorsolateral PFC of the monkeys in the active and passive tasks. **b)** The peri-stimulus time histogram averaged over all the recorded neurons of monkey ELV in the two tasks. Gray areas: the cue presentation periods. **c)** Same as (b) but for monkey ADR. **d)** Same as (b) but for monkey NIN. Neurons in NIN were recorded with varying delay lengths. Their delay periods were truncated to 1s before they were assembled as a pseudo-population.

### M.2. Neural representations, neural geometries, and representational dissimilarity vectors

We use the data of monkey ELV as an example to explain the working definitions of neural representations, neural geometries, and representational dissimilarity vectors (RDV). Although most of the 384 neurons of monkey ELV were recorded individually, we analyzed them together as if they were from the same population. We analyzed the populational data in a 384D neural state-space. Neurons' firing rate at a moment determined a point in the neural state-space. During a short time window, a stimulus stimulated a cluster of points in the neural state-space. We chose the cluster center as the neural representation of the stimulus. The neural representations (i.e., cluster centers) of different stimuli in the neural state-space together formed a geometrical structure. This structure was termed as a neural geometry. We could summarize various characteristics of a neural geometry into RDVs for representational similarity analysis. Here, we mainly used the angular RDV. An angular RDV comprised the angles between any two of the ordered arrows emitting from stimulus (neural representation) 9 to one in stimuli 1 to 8 (neural representations) (Fig. 1c–f). Accordingly, each angular RDV had $\binom{8}{2} = 28$ angular values. We also used the distance RDV. A distance RDV comprised the Euclidean distances between any two of the ordered stimuli (neural representations). Accordingly, each distance RDV had $\binom{8}{2} = 28$ distance values.

**M.3. Stimulus-neural geometry correlation**

We used correlation between the RDV of the stimulus geometry and that of the neural geometries in a sampling time window to quantify the dynamic similarity between the geometries. This was done with a sliding time window whose width was 0.2s and whose time step was 0.02s. For the angular RDVs, each RDV had 28 components. To increase the robustness of the correlation, we needed longer and richer patterns. To do so, we divided the sample window into 4 sub-windows (Fig. M2). In each sub-window, a 28-component RDV of the neural geometry were created. We then z-scored the RDV for eliminating delusive correlation caused by the magnitude difference across the sub-windows. The z-scored RDVs were concatenated to a 112-component pattern. On the other hand, the 28-component RDV of the stimulus geometry was repeated 4 times, z-scored, and concatenated to create a 112-component pattern as well. We then computed Pearson's correlation coefficient of the two concatenated patterns. This method enhanced the stability when the two patterns were uncorrelated, while ensuring a reliable reflection of actual geometry similarity. The baseline was created by randomly permutating the labeling of the stimuli 100 times. Then, RDVs of the permutated stimulus geometries were generated for computing of the baseline correlations.

For visual inspection of the similarity between the RDV of the stimulus geometry and that of the neural geometries, we sampled neural geometries in three time windows: 0.05–0.25s in the cue 1 period, 0.6–1.0s and 1.6–2.0s in the delay 1 period. We chose a shorter sampling time window in the cue 1 period because the cue 1 period was relatively short itself (0.5s). We then generated the 28-component RDVs for the three time windows. For computing static correlation in these time windows, we used the 28-component RDVs without dividing and concatenating as in Fig. M2. We also used the same method to compute dynamic correlation between distance RDVs.

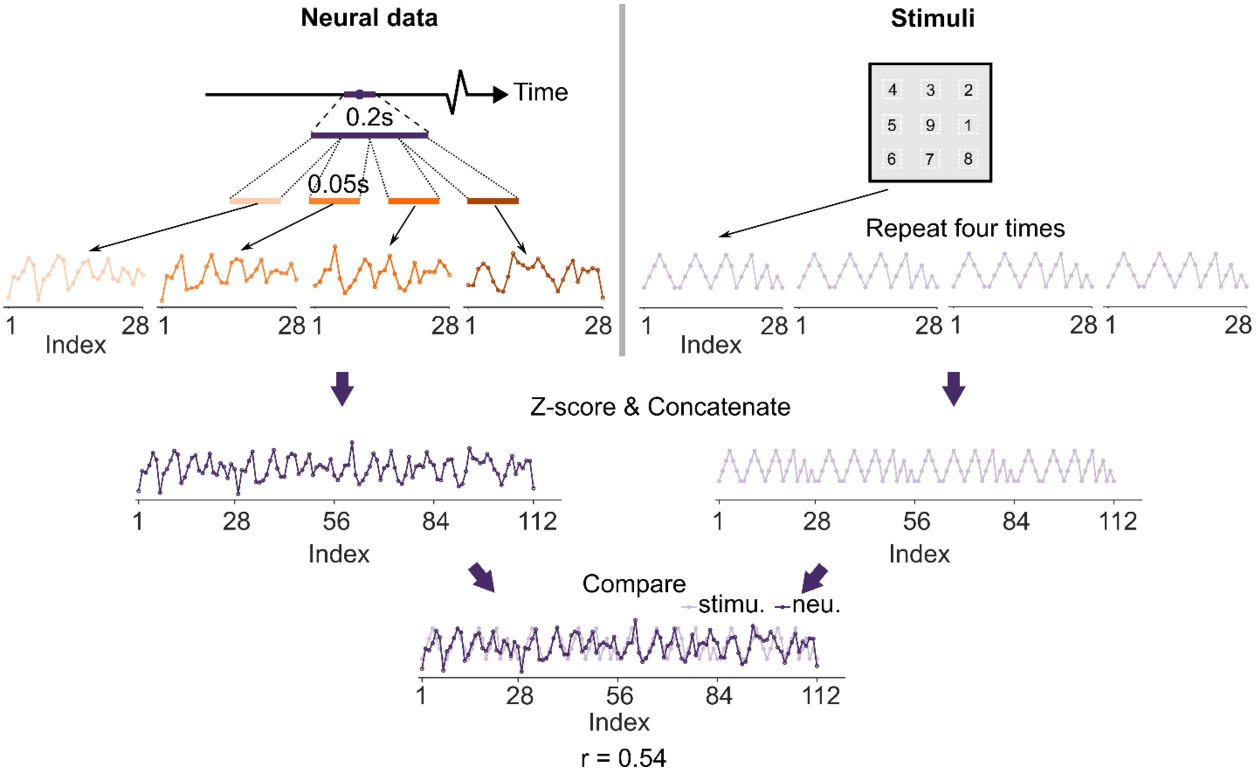

**Figure M2.** The correlation between the RDVs of the neural and stimulus geometries. To increase robustness of the correlation, four pieces of RDVs were concatenated to create a richer pattern. Pearson's correlation coefficient was then calculated for the two concatenated RDVs.

### M.4. Asymmetry measure of the neural geometries

For measuring the degree of neural geometry asymmetry, we used the standard deviation of angular RDVs, with larger standard deviation meaning higher asymmetry. Specifically, we generated the 28-component RDV of a neural geometry formed by the neural representations in a 0.4s-wide time window. The dynamic asymmetry was computed by sliding this time window with a time step of 0.02s. To create the baseline, we first generated a perfectly symmetric geometry. To do so, we initialized a 384×8 zero matrix and assigned an 8×8 identity matrix to the first 8 rows. The 8 column vectors were the locations of the vertices in the 384D space, together forming a symmetric structure. We then added Gaussian noise to each dimension of the vertex locations. The noise levels were adjusted such that the mean of the baseline matched the averaged asymmetry measure in the fixation periods. The Gaussian noise was $\mathcal{N}(0,\ 0.118)$ for the active task and $\mathcal{N}(0,\ 0.088)$ for the passive task. The symmetric geometry was randomly rotated in the 384D space to ensure its arbitrariness. We sampled 100 geometries and used their angular RDVs for computing the baseline asymmetry measure.

### M.5. Partial correlation between the sensory and mnemonic geometries

We used partial correlation to measure the similarity between the sensory geometry and the mnemonic geometry while controlling the effect of the stimulus geometry. We defined the sensory geometry as the neural geometry formed by the neural representations in the 0.05–0.25s window; the mnemonic geometry as the neural geometry formed by the neural representations in the 1.6–2.0s window. We first used the sensory geometry as the reference. We compared angular RDVs of the neural geometries across a trial against that of the reference, using the dividing and concatenating technique as in Fig. M2. Let the Pearson's correlation between the neural geometry (N) in a time window and the reference geometry (R) be $r_{NR}$; between the neural geometry and the stimulus geometry (S) $r_{NS}$; between the reference geometry and the stimulus geometry $r_{RS}$. The partial correlation between the neural geometry and the reference geometry while controlling the effect of the stimulus geometry is

$$r_{NR,S} = \frac{r_{NR} - r_{NS}r_{RS}}{\sqrt{(1 - r_{NS}^2)(1 - r_{RS}^2)}}.$$

The dynamic partial correlation was computed used a sliding window method whose width was 0.2s and time step was 0.02s. Later, we used the mnemonic geometry as the reference and computed partial correlation in the same way.

**M.6. Principal component analysis**

We applied principal component analysis (PCA) on the neural representations in a time window. Specifically, in a time window, the 8 centers of the clusters in response to the stimuli were computed as the neural representations in the 384D neural state-space. Then, the standard PCA function in Matlab was called to do the analysis.

**M.7. Visualization of neural geometries in 3D spaces**

We created the 3D spaces for visualizing neural geometries formed by 4 neural representations. We specified the 3 coordinate axes of the 3D spaces using support vector machines (SVM) as follows. Without losing generality, let the neural representations be of stimulus 2, 4, 6, and 8. We grouped the representations into two classes of equal size in three different ways: {2, 4} versus

{6, 8}, {2, 6} versus {4, 8}, and {2, 8} versus {4, 6}. We applied linear SVM to each of the classifications. All the classification accuracies were 1.0. The resulting normalized optimal separation vectors, $\hat{x}$, $\hat{y}$, and $\hat{z}$, were approximately orthogonal; the angles between them were in the range [75°, 100°]. We then forced the axes to be mutually orthogonal by applying QR decomposition. Specifically, we initialized a 384×384 identity matrix $A$. We then replaced the first 3 columns of $A$ with the optimal separation vectors. We applied the QR function in Matlab to matrix $A$: $Q$ = qr($A$). The QR function is theoretically equivalent to the Gram-Schmits method which ensures the columns of Q are mutually orthogonal, while the first columns of $Q$ stay close to the first columns of $A$ as much as possible (Strang, 2022). Thus, matrix Q defined a new 384D space. We then transformed the coordinates of the neural representations in the original neural state space to the new space by $b = aQ$, where $a$ was the 384D row vector specifying the coordinates of a neural representation in the original neural state-space, and $b$ was the 384D coordinates of the neural representation in the new space.  The row vector $b$ was truncated to the first 3D for visualizing the neural representation in the 3D space.

**M.8. Average ranked area ratio of 2D projections**

Once the neural geometries formed by 4 neural representations were visualized in a 3D space, we immediately got their 2D projections on the $x$-$y$, $y$-$z$, and $x$-$z$ planes. The looped sequence in each 2D projection created a polygon. We used the shoelace formula (Braden, 1986) to compute the area of the polygons: $s_{xy}$, $s_{yz}$, and $s_{xz}$. For instance:

$$S_{xy} = \frac{1}{2}\left(\begin{vmatrix} x_1 & x_2 \\ y_1 & y_2 \end{vmatrix} + \begin{vmatrix} x_2 & x_3 \\ y_2 & y_3 \end{vmatrix} + \begin{vmatrix} x_3 & x_4 \\ y_3 & y_4 \end{vmatrix} + \begin{vmatrix} x_4 & x_1 \\ y_4 & y_1 \end{vmatrix}\right),$$

where $(x_i, y_i)$ is the $i$th point of the $x$-$y$ projection. We ranked the areas $[s_{xy}, s_{yz}, s_{xz}]$ from in a descending order as $[s_{hi}, s_{mid}, s_{lo}]$. The ranked area ratios were

$$\left[\frac{s_{\mathrm{hi}}}{s_{\mathrm{mid}}}, \frac{s_{\mathrm{hi}}}{s_{\mathrm{lo}}}, \frac{s_{\mathrm{mid}}}{s_{\mathrm{lo}}}\right].$$

The average ranked area ratio took the mean of the ranked area ratios.

**M.9. Embedded dimensionality**

We estimated the embedded dimensionality of the 8 neural representations in the neural state-space by following the shattering dimensionality method (Bernardi et al., 2020). For 8 neural representations, theoretically there are $2^8 = 256$ possible binary classifications. Shattering dimensionality (SD) is the actual number of linearly separable binary classifications among all possibilities. We obtained SD by applying linear SVMs on each of the 256 possible binary classifications and evaluated its classification accuracy. We found all possible binary classifications were perfectly separable for any time window across a trial. That is, SD = 256. Because SD = $2^8$, we therefore concluded the embedded dimensionality was $8 - 1 = 7$, where the subtracted 1 was due to the 1 degree of freedom (Rigotti et al., 2013).

**M.10. Effective dimensionality and information entropy**

Effective dimensionality (ED) is closely related to the explained variances computed by PCA (Del Giudice, 2021; Farrell et al., 2022). For the 8 neural representations in a time window, the ranked explained variances, in descending order, were $[\lambda_1, \lambda_2, \ldots, \lambda_7]$. The effective dimensionality of the neural representations was defined as

$$\mathrm{ED} = \frac{(\sum_{i=1}^{7} \lambda_i)^2}{\sum_{i=1}^{7} \lambda_i^2} = \left(\frac{1}{\sum_{i=1}^{7} \bar{\lambda}_i^{\,2}}\right),$$

where $\bar{\lambda}_i = \lambda_i / \sum_{i=1}^{7} \lambda_i$ was the normalized explained variance. According to this formula, concentration of explained variance in a few PCs corresponds to a lower ED. In extreme, if all the explained variance concentrated to the first PC, all the neural representations were along a line in the neural state-space. In the case, the ED was 1. Conversely, a more uniform distribution of explained variance across PCs corresponds to a higher ED. In extreme, if the explained variance distribution was perfectly uniform, the neural geometry was symmetric. In this case, the ED was 7. Between the two extremes, ED was in the range (1, 7), regardless of the embedded dimensionality of the neural representations.

It has been shown in previous work (Del Giudice, 2021) that ED is closely related to Shannon's information entropy, giving it information theoretic interpretation. The interpretation, however, is different from the communicative view of entropy in information theory. In communication, entropy quantifies the amount of a stochastic event's uncertainty that can be removed by receiving the information about the true state of the event. Thus, entropy is about the ambiguity of the event to be communicated. Here, the ambiguity is about information representation in terms of a neural geometry. It is the uncertainty of one neural geometry in representing several independent pieces of information. Independent pieces of information are the PCs, commonly regarded as the feature axes. For the 8 neural representations in this study, the embedded dimensionality was 7. Thus, there were 7 feature axes $F_i$, $1 \leq i \leq 7$. The normalized explained variances along feature axes $F_i$ was $\overline{\lambda}_i$, which have the following properties

1) $\overline{\lambda}_i \geq 0$ for $1 \leq i \leq 7$;
2) the normalized explained variance of a $n$-D subspace is $\sum_{i=1}^{n} \overline{\lambda}_i$, $n \leq 7$; and
3) $\sum_{i=1}^{7} \overline{\lambda}_i = 1$.

Therefore, the normalize explained variances fulfill the three axioms of probability, thus can be interpreted as the probability of representing feature $F_i$ by the neural geometry: $p(F_i) = \overline{\lambda}_i$. We then computed Shannon's entropy

$$H = -\sum_{i=1}^{7} p(F_i)\log_2\big(p(F_i)\big) = -\sum_{i=1}^{7} \bar{\lambda}_i \log_2(\bar{\lambda}_i).$$

According to this formula, a symmetric neural geometry whose $\bar{\lambda}_i$ are uniformly distributed, entropy $H$ obtains its maximal value 2.8. An extremely skewed neural geometry, which has $\bar{\lambda}_1 = 1$ and $\bar{\lambda}_i = 0, 2 \leq i \leq 7$, has the minimal entropy 0.

**M.11. Off-plane Distance**

We developed the off-plane distance to measure the degree of flattening of a neural geometry towards the structure resembling the stimulus geometry. The idea case was that the neural geometry was flattened to 2D, same as the stimulus geometry (Fig. M3a). In this case, the neural geometry was on a flat plane. The vectors connecting the neural representations lay on this plane. That is, if we chose two nonparallel vectors and found the plane spanned by the two vectors, we knew the other vectors were in the plane (Fig. M3b). However, our results show the neural geometries remained high-dimensional throughout a trial (Fig. 3c), but its projection to the first 2 PCs obtained by PCA were in the similar structure as the stimulus geometry (Fig. 3a), as depicted in Fig. M3c. To accommodate these findings, we restricted the vectors connecting neural representations to be the diametric vectors with no common end. Any two diametric vectors spanned a flat plane. Flatness was redefined as the other diametric vectors being parallel to the plane (Fig. M3c). To quantify flatness, we first randomly chose two diametric vectors (Fig. M3d). We then found the plane spanned by the two vectors. Lastly, for each remaining diametric vector, we computed the distance residual between the vector and its projection into the found plane. This distance residual was named off-plane distance. A higher averaged off-plane distance means a less flat neural geometry.

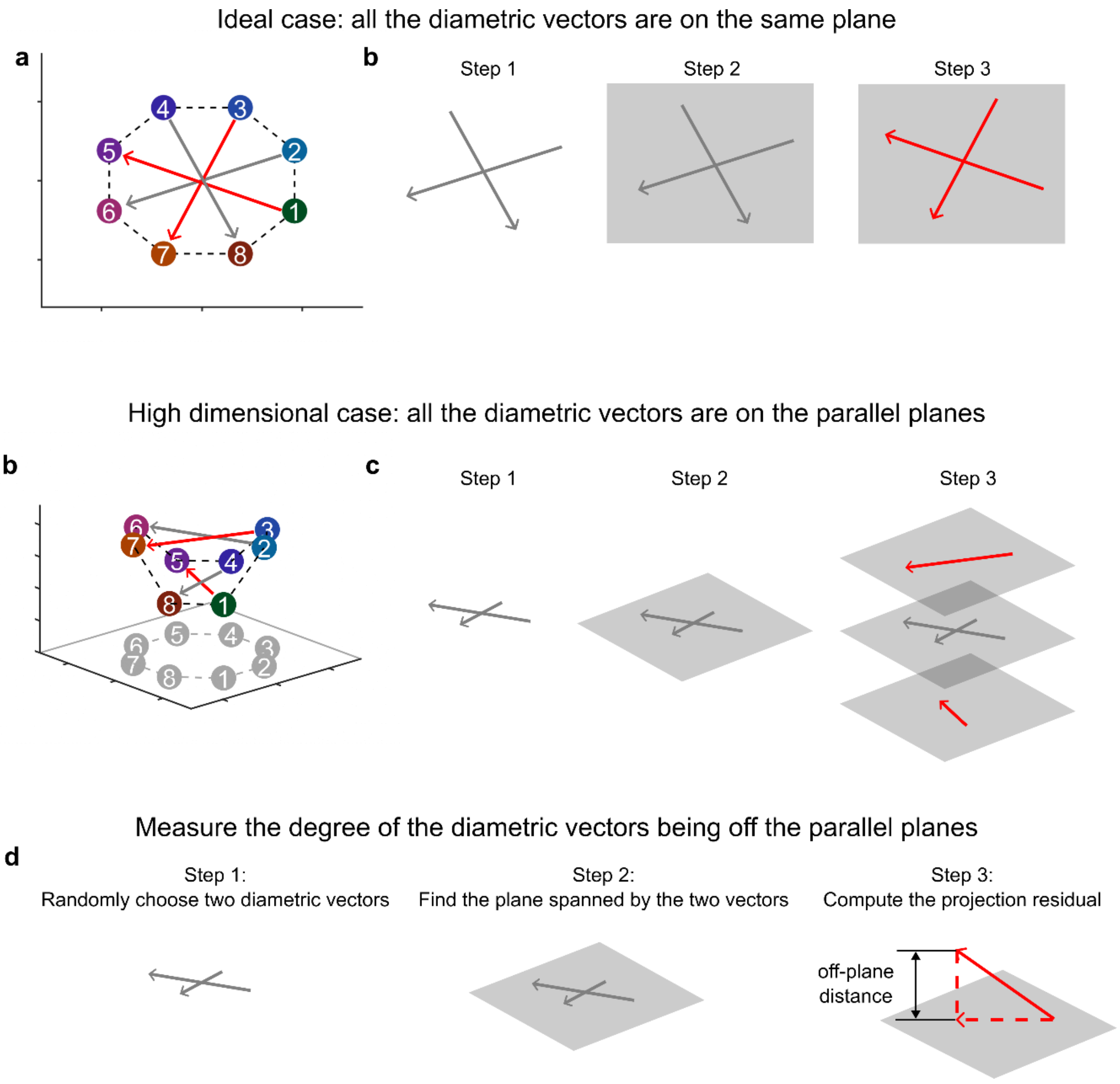


**Figure M3.** Explanation of the off-plane distance. **a)** The idea case in which the neural geometry is of the same dimensionality as the stimulus geometry. **b)** For the idea case in (a), any two chosen non-parallel vectors spanned a flat plane (steps 1–2). The remaining vectors lay in the plane (step 3). **c)** In the actual case, the neural geometry was high-dimensional while its projection to the first 2 PCs was similar to the stimulus geometry. **d)** For the actual case in (b), any two chosen diametric vectors spanned a flat plane (steps 1–2). A flat neural geometry had other diametric vectors parallel to the plane (step 3). **d)** The procedure of computing off-plane distance. Step 1: randomly choose two diametric vectors. Step 2: find the plane spanned by the two vectors. Step 3: compute the projection residual of other diametric vectors to the plane as the off-plane distance.

### Supplementary Notes and Figures

In this Supplementary Notes and Figures, we present additional results in support of the results and conclusions in the main text.

**N.1. Decoding analysis**

The visuospatial delayed-match/nonmatch task has two delay periods. From the characteristic of the task, we speculated that the first delay period was for temporarily holding the stimulus information of the first cue, and the second delay period was for making and holding a decision before responding. We applied decoding analysis to verify the speculation. We tested the generalization accuracy of the stimulus identity and the match/nonmatch decision across a trial. If the speculation was true, we expected to see the successful decoding of the stimulus identity in the first delay period and of the match/nonmatch decision in the second delay period, but not the other way around.

For decoding the stimulus identity information before cue 2 onset, we randomly split the experiment trials into two equal sized sets, and took trial averages for each set. We used one set for training a linear SVM (default parameters in Matlab) and the other for testing the generalization accuracy. The SVM was to classify the diametrically opposite stimuli, for instance, stimulus 1 versus stimulus 5. There were in total 4 pairs of opposite stimuli. We then pooled the generalization accuracies together for computing the mean and standard deviation. For the decoding analysis after cue 2 onset, where there were match and nonmatch trial depending on the consistency of cue 1 and cue 2, we did the training-testing split for both kinds of trials, and pooled the generalization accuracies together. These analyses were done for both the active and passive tasks, where results of the passive task were used as the baseline. The dynamic accuracies were obtained by using a sliding window of 0.2s-wide. The time step was 0.02s.

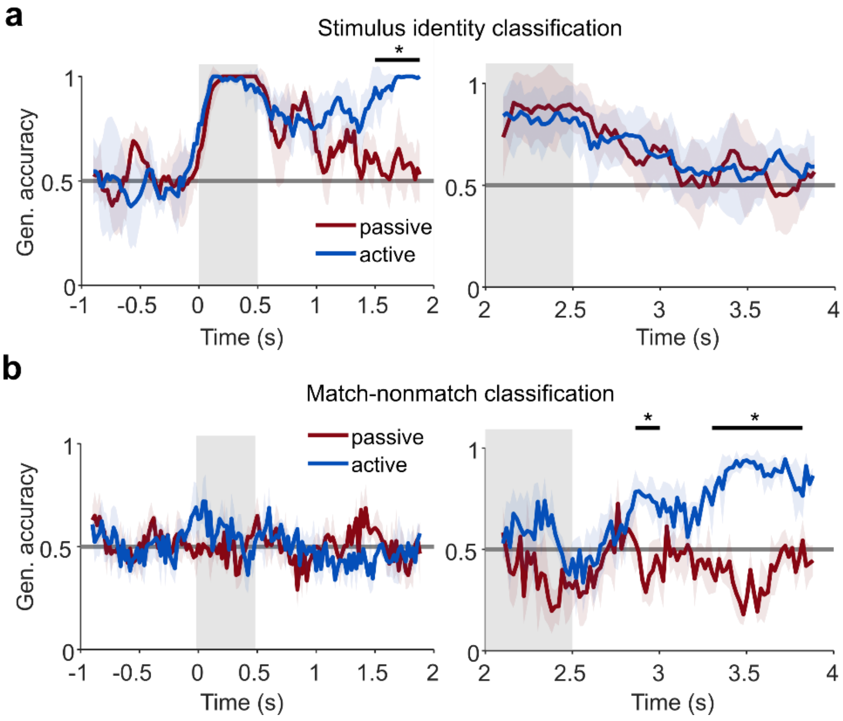


**Figure S1**. The decoding analysis for distinguishing stimulus identities and match/nonmatch decisions. **a)** The generalization accuracies of stimulus identities in both the active and passive tasks, and before (left) and after (right) cue 2 onset. Gray line: the chance level **b)** Same as (a) but for the generalization accuracies of match/nonmatch decisions. Asterisk indicates significance $p < 0.05$.

The generalization accuracies of the active and passive tasks were similar initially: around the chance value in the fixation period and jumping to perfect accuracy in the cue 1 period (Fig. S1a, left). Then, the generalization accuracies forked during the delay 1 period: the accuracy of the passive task data decreased gradually to the chance value, while the accuracy of the active task data maintained a moderate value and then resurged in the late delay, with the difference between the accuracies being significant (t-test, $p < 0.05$). This result is consistent with previous findings (Meyers et al., 2012). The discrepancy of generalization accuracies was as expected, because in the active task the monkey must hold the stimulus identity information, whereas the information was not necessary for the passive task.

However, entering the cue 2 period, despite the initial high accuracies driven by the stimulus presentation, the generalization accuracies of both the active and passive tasks gradually decreased to the chance level during the delay 2 period (Fig. S1a, right). This result verified that stimulus identity became task-irrelevant in the delay 2 period. Instead, the delay 2 period was speculated as the time when match-nonmatch decisions were made and held.

For decoding the match-nonmatch decisions, we took the trial averages of firing rates for the match trials and nonmatch trials, respectively. We then randomly drew neural representations of six stimuli for training, and the neural representations of the remaining two stimuli were reserved for testing. We repeated the training-testing procedure 4 times. The generalization accuracies were pooled for computing the mean and standard deviation. The dynamic accuracies were obtained by using a sliding window of 0.2s-wide. The time step was 0.02s.

Before the onset of cue 2, the match-nonmatch information must not be decodable. Indeed, the generalization accuracies fluctuated around the chance value for both the active and passive tasks (Fig. S1b, left). In contrast, the generalization accuracies gradually ramped up during the delay 2 period in the active task, significantly greater than the accuracies in the passive task (t-test, $p < 0.05$), which still fluctuated near the change level (Fig. S1b, right). Taken together, the results verified our speculation that the first delay period was for memorizing the stimulus information and the second delay period was for making and maintaining match/nonmatch decisions.

**N.2. Correlation and asymmetry of distance RDVs**

Beside the angular representational dissimilarity vectors (RDV), we also used distance RDVs as a description of the stimulus and neural geometries. Specifically, for the stimulus geometry of 8 locations, we computed the Euclidean distances between each pair of the stimuli, and assembled the $\binom{8}{2} = 28$ distances into a RDV vector. Similarly, we obtained the distance RDVs of the neural geometry in the neural state-space. As in Fig. 2, we analyzed the correlation between the distance RDV of the stimulus geometry and that of the neural geometry for both the active and passive data. Also, we analyzed the standard deviation of the distance RDVs as a measure of the neural geometry asymmetry. The RDV standard deviation was normalized by the mean of the distance RDVs to avoid the influence of changing neural geometry size across time:

$$\text{Scaled RDV std.} = \frac{\text{std}(\ RDV\ )}{\text{mean}(\ RDV\ )}.$$

The dynamic time courses were obtained using a sliding window across the trial (width: 0.2s for correlation, 0.4s for RDV standard deviation; time step: 0.02s).

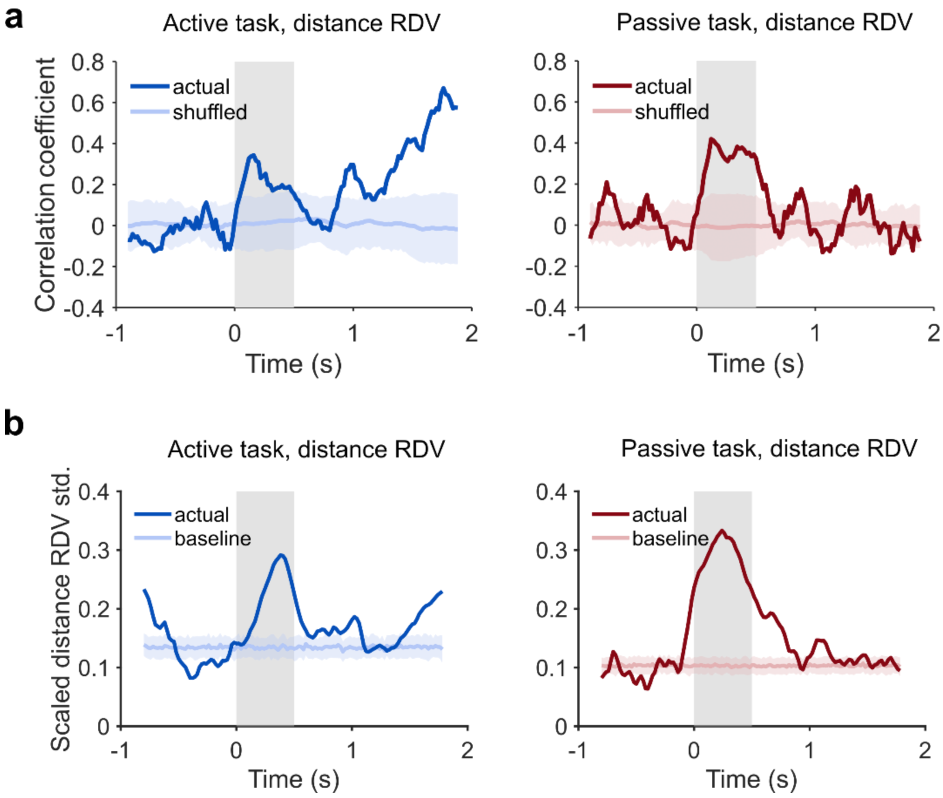


**Figure S2.** Correlation and asymmetry of distance RDVs. **a)** The Pearson's correlation between the distance RDV of the stimulus geometry and that of the neural geometry across the trial in the active task (left) and the passive task (right). Gray region indicates the cue 1 presentation period. The baseline was created by randomly shuffling the labels of the stimuli 100 times, thus shuffling the components of the distance RDV. The mean (light colored line) and standard deviation (light shaded area) are shown. During the delay 1 period, the correlation in the active task decreased first and then increased, while the correlation in the passive task decreased and fluctuated near 0 correlation. **(b)** The scaled distance RDV standard deviation of the neural geometry across the trial, in the active task (left) and the passive task (right). Gray region indicates the cue 1 presentation period. The baseline was created by sampling 100 geometries from a space generated by adding Gaussian noise to the perfectly symmetric geometry. The noise levels were adjusted by matching the mean of the baseline to that of the data in the fixation period (-1s– 0s). The mean (light colored line) and standard deviation (light shaded area) are shown. Similar to the correlation, the scaled distance RDV standard deviation in the active task decreased first and then increased in the delay 1 period. The scaled distance RDV standard deviation in the passive task decreased and remained near the baseline in the same period. These results are consistent with the analysis of the angular RDVs in Fig. 2.

#### N.3. Stimulus-neural geometry correlation in the delay 2 period

Neural activities in the delay 2 period provided functions different from the activities in the delay 1 period. Activities in the delay 2 period were for making and holding decisions. Therefore, the match/nonmatch decisions were task-relevant in this period, and the stimulus information became irrelevant (Fig. S1). As a result, the stimulus relation would be not represented in the delay 2 period. Accordingly, the correlation between the stimulus geometry and the neural geometries in the delay 2 period must be low. We tested this prediction by applying the same method as in computing the correlation in the delay 1 period (see M.3. Stimulus-neural geometry correlation). Dynamic correlation was obtained with a sliding time window whose width was 0.2s and time step was 0.02s. The results are in Fig. S3. The stimulus-neural geometry correlation indeed decreased to near 0 in the delay 2 period for both the active and passive task data, indicating that when the stimulus relation was not task-relevant anymore, the neural geometries ceased transforming toward the stimulus geometry.

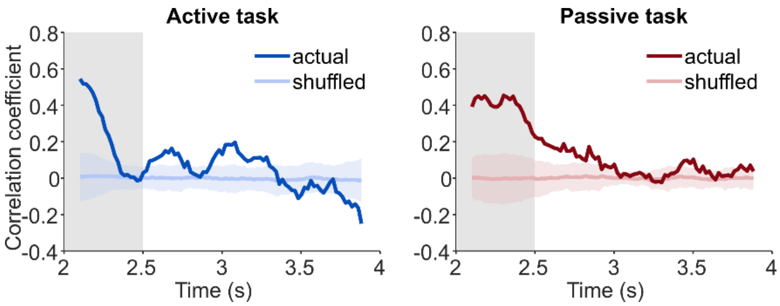


**Figure S3.** Stimulus-neural geometry correlation in the delay 2 period. **a)** The dynamic Pearson's correlation between the angular RDV of the stimulus geometry and that of the neural geometry in a sliding time window across the cue 2 and delay 2 periods of the active task. The correlation decreased to near 0 in the delay 2 period. The baseline was created by randomly shuffling the labels of the stimuli 100 times. The mean (light colored line) and standard deviation (light shaded area) are shown. Gray region: the cue 2 period. **b)** Same as (a) but for the passive task data. Similarly, the correlation decreased to 0. The correlation was less oscillated in the delay 2 period of the passive task.

#### N.4. PCA of the passive task data

We applied PCA to the neural representations in three sampling time windows of the passive task. The three time windows were in the cue 1 period (0.05s–0.25s), early delay 1 period (0.6–1.0s), and late delay 1 period (1.6s–2.0s). Since in the passive task, the monkeys did not need to maintain stimulus information during the delay, we expected the neural representations to reflect the structure of the stimulus geometry only in the cue 1 period. The result in Fig. S4 show it is indeed this case. This result provides a control for the detected stimulus relation in the late delay 1 period of the active task (Fig. 3a).

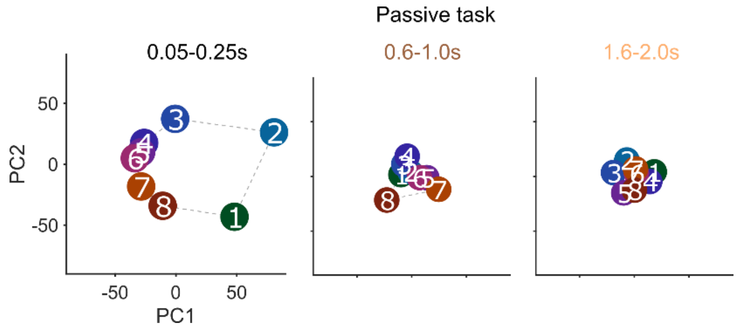


**Figure S4.** Projections of the passive task data to the first two PCs. Only in the cue 1 period did the projections of the neural representations displaying the structure of the stimulus geometry (left). This structure did not show up in the projections during the early delay 1 period (middle), and the late delay 1 period (right).

### N.5. Partial correlation analysis

Partial correlation analysis on the angular RDVs in the active task revealed that the neural geometries across the trial were dynamic. Here, we used the same method to analyze the dynamic partial correlation between the angular RDVs in the passive task (Fig. S5a), and between the distance RDVs in the active and passive tasks (Fig. S5b).

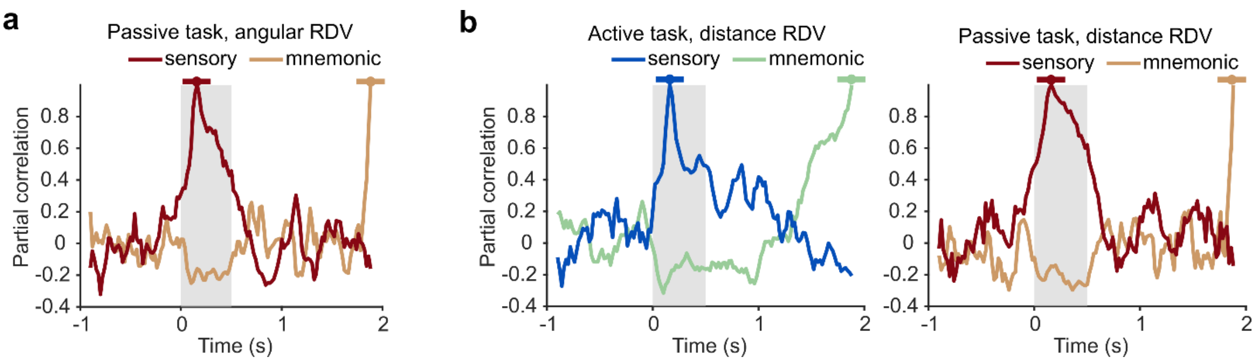


**Figure S5.** Partial correlation between neural geometries. **a)** The dynamic partial correlation between the angular RDVs in a sliding window across the trial and that of the sensory and mnemonic geometries, respectively, in the passive task. During the delay 1 period, the partial correlation to the mnemonic geometry oscillated around 0, indicating uncorrelated neural geometries. Sliding window: width = 0.2s, time step = 0.02s. **b)** The dynamic partial correlation between the distance RDVs in a sliding window and that of the sensory and mnemonic geometries, respectively, in the active task (left) and in the passive task (right). In the active task, the result is consistent with Fig. 3b, where the partial correlation to the mnemonic geometry gradually increased in the delay 1 period, but the partial correlation between the sensory geometry and the mnemonic geometry was low. In the passive task, the result is consistent with (a), where the partial correlation to the mnemonic geometry oscillated around 0.

### N.6. Permutation analysis of relational representations

To test the robustness of correlation as a measure of relation representation, we permuted the sequence of the stimuli (i.e., 1-2-3-4-5-6-7-8). Here we also recorded the number of inversions of permutated sequences with respect to the original sequence. An inversion is the reversed ordered of a pair of digits. For instance, there is a pair 1-3 in the original sequence, regardless of how many digits between 1 and 3. A permutated sequence having 3-1, regardless how many digits in between, has one additional inversion. We aimed to compare the original and permutated sequences to find all inversions.

Before comparison, some adjustment must be applied to the permutated sequences for an accurate number of inversions. Because the sequence of the stimuli is looped, that is, 1-2-3-4-5-6-7-8 is the same as 8-1-2-3-4-5-6-7, we always aligned the original and permutated sequences such that the digit 1 was always the first digit in the sequences. Also, because the looped sequence is symmetric, that is, the original sequence 1-2-3-4-5-6-7-8 and the flipped original sequence 1-8-7-6-5-4-3-2 represent the same spatial relation, we compared the permutated sequence to both the original and the flipped original sequences, and found the minimal number of inversions as the result of the comparison. According to this procedure, the exemplar sequence 1-2-3-4-5-7-6-8 had one inversion and 1-7-2-3-4-5-6-8 had five inversions. More inversions indicated higher dissimilarity between two sequences. We segregated the permutated sequences according to their inversions and computed the correlations between the angular RDVs derived from these sequences and the neural geometry in the late delay 1 period (1.8–2.0s). The results are in Fig. S6. This control analysis strengthens the conclusion that what was established in the neural state-space was correct neural representations of the correct stimulus relation in the late delay 1 period for the active task.

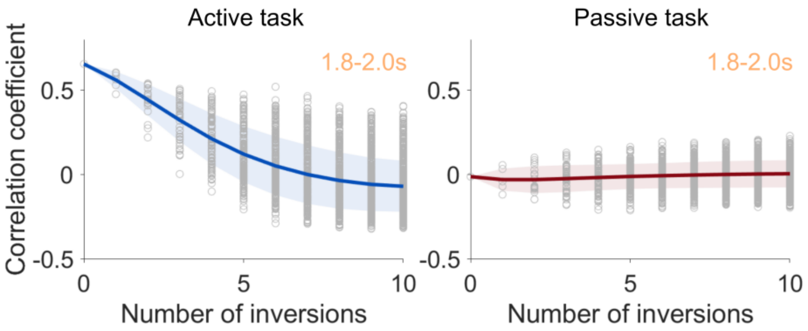


**Figure S6.** Permutation analysis of the relational representations. **a)** The correlation between the angular RDV of the neural geometry in the late delay 1 period and that of the permutated stimulus geometry for the active task data. The number of inversions quantifies the dissimilarity between the actual stimulus geometry and the permutated stimulus geometry. The Pearson's correlation coefficient decreases across the number of inversions. Solid line: mean correlation, shaded area: standard deviation, gray circles: data points. **b)** Same as (a) but for the passive task data. The mean correlation coefficient remains at 0 for any permutation.

#### N.7. Inter-representation distance analysis

To further verify the changing asymmetry of the neural geometries, we computed Euclidean distances between each pair of representations in the high-dimensional neural state-space in three sampling time windows within the cue 1 period (0.05s–0.25s), early delay 1 period (0.8s–1.0s), and late delay 1 period (1.8s–2.0s), respectively. If a neural geometry is fully symmetric, we would observe uniform inter-representation distances. Otherwise, if a neural geometry is asymmetric towards reflecting the stimulus relation, we would expect that the distribution of the inter-representation distances reflected characteristics of the stimulus geometry. One characteristic is the distance between the spatial locations. For instance, stimulus 2 is closer to stimulus 1 than stimulus 5. For simplicity, we used Manhattan distances to quantify the difference. Manhattan distances count the number of links between the two stimuli in the looped sequence of the stimulus geometry. For instance, in the looped sequence 1-2-3-4-5-6-7-8-1, the Manhattan distance between stimulus 1 and stimulus 5 is $m = 4$. Asymmetry neural geometries reflecting the stimulus relation would result in larger inter-representation distances for larger $m$. We examined the inter-representation distances in the three sampling time windows. Results are shown in Fig. S7. The inter-representation distances show clear evidence of asymmetry reflecting the stimulus relation in the late delay 1 period in the active task and in the cue 1 period in the passive task.

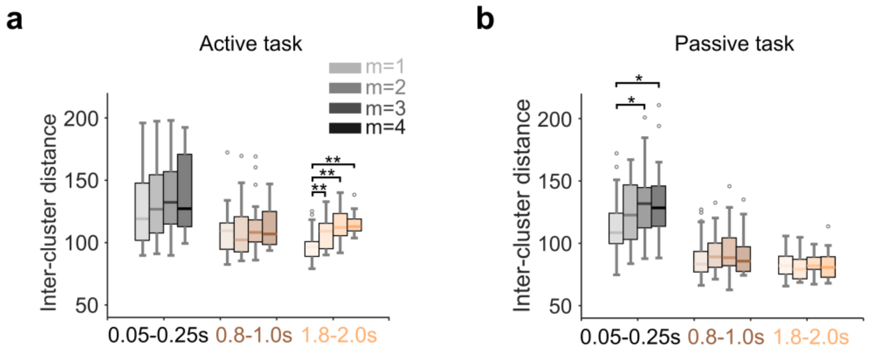


**Figure S7.** Box plots of the inter-representation distances across three sampling time windows. **a)** For the active task data, ANOVA revealed significant difference existed only in the late delay 1 period ($F(3, 108) = 13.95$, $p < 0.01$, right). In this time window, the medians were monotonically increasing with respect to m, as expected. Pairwise post-hoc comparisons showed that inter-representation distances at $m = 2$, 3 and 4 were significantly larger ($p < 0.01$) than the distances at $m = 1$. However, the inter-representation distances in the early delay 1 period were at the same level, indicating a symmetric geometry ($F(3, 108) = 0.52$, $p = 0.671$, middle). The inter-representation distances in the cue 1 period had an increasing trend with respect to m, as expected, although the difference was not significant

$(F(3, 108) = 1.32, p = 0.273$, left). Asterisk: statistical significance $p < 0.01$. Gray circles: Outliers. **b)** For the passive task data, the median distances were almost equal and no significant difference was detected in the early and late delay 1 periods ($F(3, 108) = 0.45$, $p = 0.716$, right, and $F(3, 108) = 0.32$, $p = 0.814$, middle). However, the inter-representation distances in the cue 1 period were significantly different ($F(3, 108) = 3.84$, $p = 0.012$, left). Pairwise post-hoc comparisons showed that inter-representation distances at $m = 3$ and 4 were significantly larger ($p < 0.01$) than the distances at $m = 1$, showing an increasing trend.

### N.8. Three-D visualization of the neural geometries in the passive task

In comparison to the visualization of neural geometries in the active task, we also the same method (i.e., SVM-based visualization, see Supplementary Methods) to visualize the neural geometries in the passive task. Since the monkeys did not need to memorize the stimulus information for completing the task, we expected the neural geometries in the delay 1 period to be symmetric. The results are shown in Fig. S8. Indeed, the neural geometries were less asymmetric in the delay 1 period.

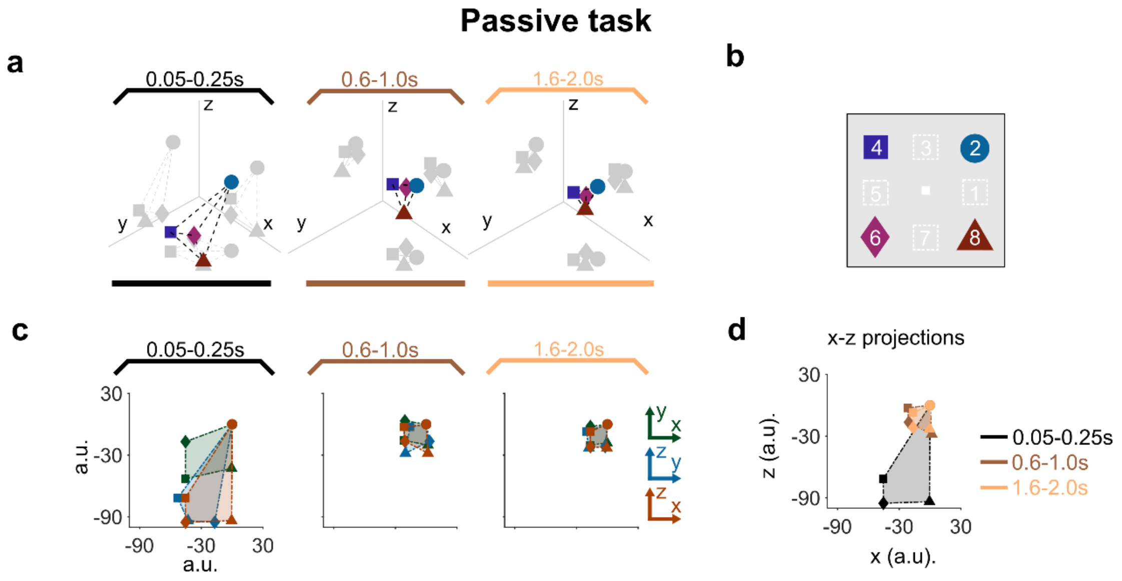

**Figure S8.** Visualization of the neural geometries in the passive task. a) Visualization in 3D spaces of the neural geometries formed by representations of the stimuli in (b) and in the three sampled time windows in the cue 1, early delay 1, and late delay 1 periods, respectively. Marker shapes refer to stimulus identities. **b)** The stimulus geometry with four selected spatial stimuli. **c)** The 2D projections of the neural geometries in (a), grouped according to the sampling time windows. According to the 2D projections, the neural geometry in the cue 1 period was more elongated (average ranked area ratio = 1.31) than the neural geometries in the early delay 1 period (average ranked area ratio = 1.08) and in the late delay 1 period (average ranked area ratio = 1.1). As a result, visually the neural geometries in the delay 1 period are more symmetric. **d)** The x-z projections in the three sampling time windows. The 2D projections in the delay 1 period were similar.

### N.9. Change of information entropy over the relation reconstruction

Over the relation reconstruction in the delay 1 period of the active task, the effective dimensionality decreased (Fig. 4b), meaning dimensionality compression was in action. From the Supplementary Method section (M.10. Effective dimensionality and information entropy), we find effective dimensionality is related to information entropy through the normalized explained variances $\bar{\lambda}_i, i = 1, \dots, 7$. We therefore computed the information entropy for the active and passive task data in the mid-late delay 1 period, where the relation reconstruction happened. The dynamic entropy used a sliding time window method (width: 0.2s, time step:0.02s). The results are in Fig. S9. We found the information entropy in the active task monotonically decreased over this period. The decreasing information entropy gave a clear interpretation: the neural geometry became increasingly certain in representing the information of the stimulus relation.

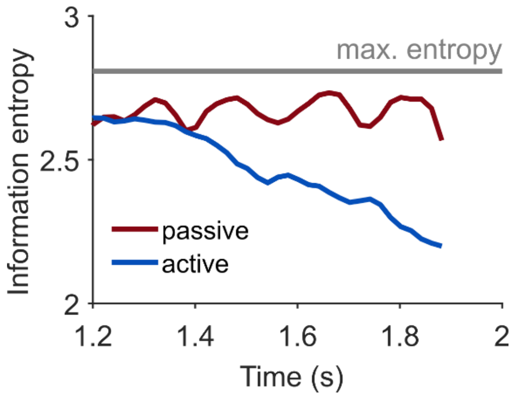


**Figure S9.** Change of information entropy over the relation reconstruction. In the active task, the information entropy monotonically decreased over the mid-late delay 1 period, showing the neural geometry became progressively certain in representing the stimulus information. In contrast, the information entropy in the mid-late delay 1 period of the passive task oscillated around a constant value. The maximal entropy was computed by assuming $\bar{\lambda}_1 = \bar{\lambda}_2 = \cdots = \bar{\lambda}_7$. The geometry in this case is perfectly symmetric. The information entropy in the passive task remained near the maximal value.

**N.10. Shattering dimensionality of the neural representations**

To estimate the embedded dimensionality of the neural representations of the 8 stimuli, we adopted the shattering dimensionality method from (Bernardi et al., 2020). This method computes the number of the actual binary classifications that are linearly separable, and compares this number with the theoretical number of all possible dichotomies. We found the actual number of linearly separable dichotomies was 256, which equaled the theoretical number $2^8$, throughout the experiment trial for both the active and passive task data. The result is shown in Fig. S10. This result shows the embedded dimensionality of the neural geometry was always 7D, the highest embedded dimensionality for 8 neural representations.

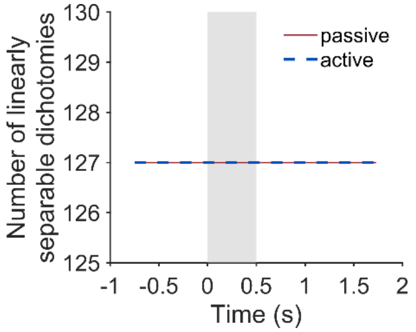


**Figure S10.** The number of unique nontrivial actual linearly separable dichotomies in both the active and passive tasks. There is one unique trivial dichotomy that is by default linearly separable: {1, 2, 3, 4, 5, 6, 7, 8} versus $\emptyset$. Also, dichotomies are symmetric, for instance, {1, 2, 3, 4} versus {5, 6, 7, 8} is the same dichotomy as {5, 6, 7, 8} versus {1, 2, 3, 4}. Therefore, the total actual linearly separable dichotomies was $(127 + 1) \times 2 = 256$, which equaled the theoretical number of all possible dichotomies $2^8$. This number was constant in the fixation, cue 1, and delay 1 periods, indicating the neural geometry always took the highest dimensionality 7 afforded by 8 neural representations.

### N.11. Complete time courses of effective dimensionality and off-plane distance

In Fig. 4b–c, the time courses of the effective dimensionality and the off-plane distance is shown for the duration of relation reconstruction (1.2s–2.0s). Here we show the complete time courses from the fixation period to the delay 1 period. During the cue 1 period, since the neural geometries were driven by the stimuli, we expected them to be flattened, and therefore the effective dimensionality and the off-plane distance would decrease in both the active and passive tasks. We used the same method as in computing the effective dimensionality and off-plane distance in Fig. 4b–c (see Supplementary Methods M10 and M11). The result is shown in Fig. S11. Indeed, both the effective dimensionality and off-plane distance were lower in the cue 1 period in the active and passive tasks, respectively. However, they quickly recovered to higher values in the early delay 1 period. Then, in the mid-late delay 1 period, they gradually decreased in the active task only. This control analysis shows that the effective dimensionality and off-plane distance could reflect the encoding of the stimulus relation.

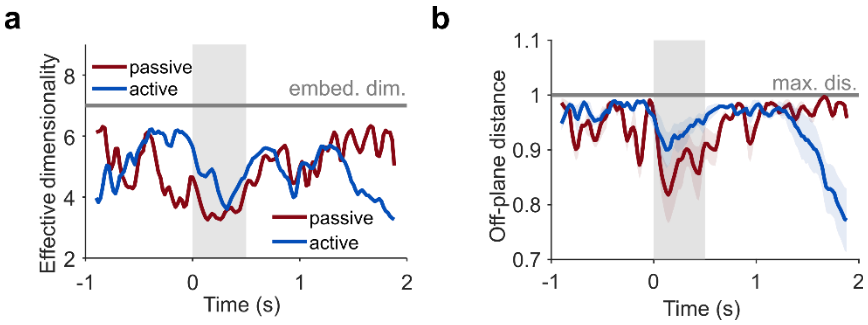


**Figure S11.** Complete time courses of effective dimensionality and off-plane distance. a) The time course of the effective dimensionality in the fixation, cue 1, and delay 1 periods. From the fixation period to the cue 1 period, the effective dimensionality decreased because of the stimulus presentation, in both the active and passive tasks. However, even before cue offset, the effective dimensionality started to recover to the similar value as in the fixation period, in the active and passive tasks. Only in the active task, the effective dimensionality fell again in the mid-late delay period, an indication of geometry flattening. Gray region: the cue 1 period. Gray line: the embedded dimensionality of the neural geometry. b) The time course of the off-plane distance in the fixation, cue 1, and delay 1 periods. The temporal change was similar to that of the effective dimensionality, showing the evidence of the relation encoding (in the cue 1 period) and relation reconstruction (in the mid-late delay 1 period). Gray line: the upper limit of off-plane distances.

**N.12. Stable subspace identification according to** (Murray et al., 2017)

Murray et al. (2017) found there exists a stable subspace for maintaining memoranda in the dynamic coding of working memory. Here we applied the same method on the data in this work, aiming to identify a stable subspace. According to  (Murray et al., 2017), we took average of the neural activities in the delay 1 period (0.75s–1.75s, the 0.25s gaps were for isolating the chosen segment). The averaging removed the time information and gave 8 neural representations as we did for short time windows. We then applied PCA on the 8 neural representations and visualized their projections in the first two PCs. The result is in Fig. S12. In the visuospatial delayed-match/nonmatch task, this procedure did not find a stable subspace in which the structure of the stimulus geometry was evident.

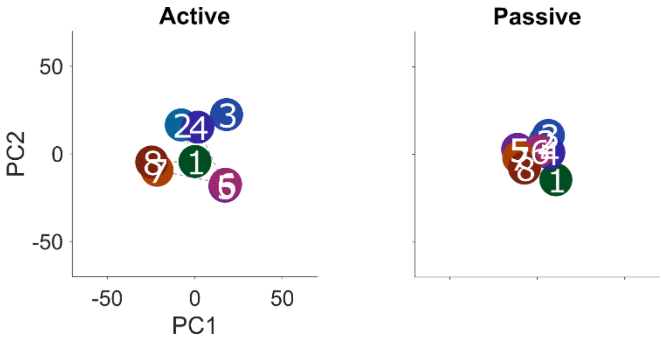


**Figure S12.** Projections of the neural representations to the first two PCs according to (Murray et al., 2017**).** The structure of a looped sequence of the representations, as in the stimulus geometry, was not apparent in the active (left) and passive (right) tasks. The procedure in (Murray et al., 2017)**,** when applied to the data of a visuospatial delayed-match/nonmatch task, failed in identifying a stable subspace in which the structure of the stimulus geometry was evident.

**N.13. Sensory subspace hypothesis**

We hypothesized that the stimulus identity was encoded in the same subspace for both cue 1 and cue 2, a sensory subspace. The implication is that the linear classifiers trained in the cue 1 period would generalize to the data in the cue 2 period. We tested this implication using linear SVM. Specifically, to extend analysis to the cue 2 period, we selected only the trials in the match condition of the experiment, and took the trial average as neurons' firing rate. We then trained linear SVMs to classify the diametric stimuli, e.g., {1} versus {5}, {2} versus {6}, etc., with the data from a 0.2s-wide time window in cue 1 period (0.05s–0.25s). We tested the classifiers on the same stimuli pair but with the data in a 0.2s-wide time window in other periods, each period was 0.5s-wide, e.g., the early fixation period (-1s–0.5s). The testing accuracies in each period in the active and passive tasks are in Fig. S13. The generalization was only good in the cue 2 period, supporting the sensory subspace hypothesis. It also indicates that this sensory subspace was orthogonal to other subspaces.

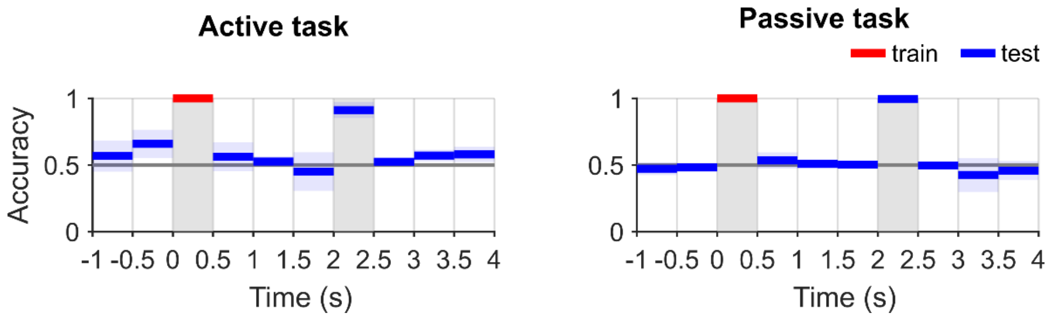


**Figure S13.** Cross-temporal generalization across the trial in the active and passive tasks. Left: linear SVM classifiers of diametric stimuli, e.g., {1} versus {5}, {2} versus {6}, etc., were trained in a 0.2s-wide time window in the cue 1 period of the active task. The classifiers were tested on the data in a 0.2s-wide time window in other periods, each period was 0.5s-wide. The temporal gaps were for isolating the time windows. The testing accuracies in the other periods except the cue 2 period were near the chance level (0.5), but the testing accuracy in the cue 2 period was close to 1, showing the classifiers generalized well to the data in the cue 2 period. Right: same as in the left panel but for the passive task data. The generalization of the classifiers trained in the cue 1 period was only good in the cue 2 period. Taken together, the result supports the hypothesis that the stimulus identity information was encoded in the same sensory subspace which is orthogonal to other subspaces.

#### N.14. Similar results in the other monkeys

The main results in this work include 1) the similarity measure in terms of the correlation between the stimulus and neural geometries, 2) the asymmetry measure in terms of the angular RDVs of the neural geometries, 3) the dimensionality measure in terms of the effective dimensionality, and 4) the targeted flatness in terms of the off-plane distance. These results were from the analysis of monkey ELV's data, because this monkey had the highest number of neurons being recorded in the active task (384 neurons) and the passive task (384 neurons) (Fig. M1). The other two monkeys provided less qualified neuron recordings (monkey ADR, the active task, 137 neurons, the passive task, 65 neurons; monkey NIN, the active task, 141 neurons, the passive task, 112 neurons). Nevertheless, we analyzed the data of monkey ADR and monkey ELV and obtained the four main results for the two monkeys. The results are shown in Fig. S14. These results reproduced the main trends observed in the results of monkey ELV.

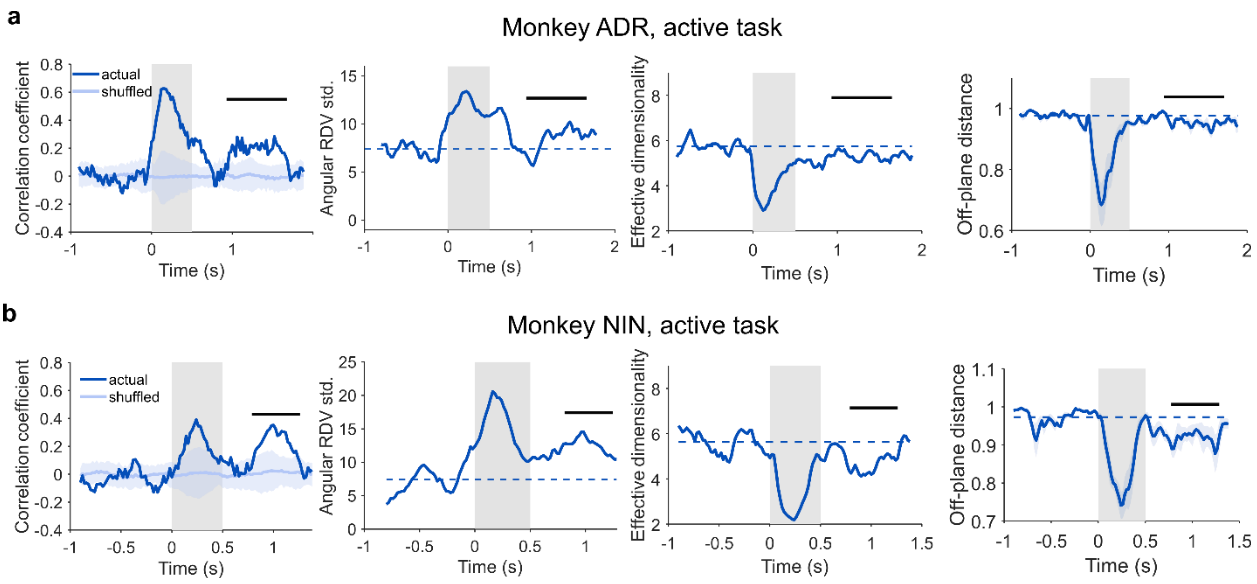


**Figure S14.** The four main results observed in the data from monkey ADR and monkey NIN. **a)** Of monkey ADR, from left to right, the time courses of the stimulus-neural geometry correlation, the angular RDV standard deviation, the effective dimensionality, and the off-plane distance in the active task. The gray region shows the cue 1 period. Importantly, the stimulus-neural geometry correlation increased in the delay 1 period. The baseline was obtained by shuffling the labels of the stimuli 100 times. The baseline mean (light colored line) and the standard deviation (light shaded area) are shown. The interval where the correlation was great than the upper bound of the baseline (mean + std.) was marked by the black bar. In this interval, the angular RDV standard deviation increased as well, indicating more asymmetric neural geometries in this interval. The dashed line acts as a reference, showing the averaged value in the fixation period. The effective dimensionality and the off-plane distance were slightly below the reference line (dashed line) in the same interval, showing targeted flattening was applied to the neural geometries in this interval. These trends of the time courses were consistent with the results of monkey ELV, although they were less obvious maybe due to the smaller number of neurons being analyzed. **b)** Same as (a) but for monkey NIN. Again, the time courses were consistent with the results of monkey ELV in the increased stimulus-neural geometry correlation and angular RDV standard deviation, and the decreased effective dimensionality and off-plane distance in the delay 1 period.
